# In a nutshell: pistachio genome and kernel development

**DOI:** 10.1101/2024.06.24.600444

**Authors:** Jaclyn A. Adaskaveg, Chaehee Lee, Yiduo Wei, Fangyi Wang, Filipa S. Grilo, Saskia D. Mesquida-Pesci, Matthew Davis, Selina C. Wang, Giulia Marino, Louise Ferguson, Patrick J Brown, Georgia Drakakaki, Adela Mena-Morales, Annalisa Marchese, Antonio Giovino, Esaú Martínez, Francesco Paolo Marra, Lourdes Marchante Cuevas, Luigi Cattivelli, Paolo Bagnaresi, Pablo Carbonell-Bejerano, Grey Monroe, Barbara Blanco-Ulate

**Affiliations:** Department of Plant Sciences, University of California, Davis, California, USA; Corto Olive, Lordi, California, USA; Department of Food Science and Technology, University of California Davis, Davis, California, USA; Regional Institute of Agri-Food and Forestry Research and Development of Castilla-La Mancha (IRIAF), IVICAM. Ctra. Toledo-Albacete s/n, 13700 Tomelloso (Ciudad Real), Spain; Department of Agricultural, Food and Forest Sciences, University of Palermo, Viale delle Scienze - Ed. 4, 90128 Palermo, Italy; CREA for Agricultural Research and Economics (CREA), Research Centre for Plant Protection and Certification (CREA-DC), Viale delle Scienze, 90128 Palermo, Italy; CREA Research Centre for Genomics and Bioinformatics, Fiorenzuola d’Arda, 29017, Italy; Instituto de Ciencias de la Vid y del Vino, ICVV, for Grape and Wine Sciences ICVV, CSIC - Universidad de La Rioja - Gobierno de La Rioja, 26007 Logroño, Spain

## Abstract

Pistachio is a sustainable nut crop with exceptional climate resilience and nutritional value. To advance pistachios as a future food source and a model system for hard-shelled fruits, we generated a chromosome-scale reference genome of the most widely grown pistachio cultivar (*Pistacia vera* ‘Kerman’) and a spatiotemporal developmental study of the hull, shell, and kernel. Our study defined four distinct stages of pistachio growth and maturation by integrating tissue-level physiological and molecular data from thousands of nuts across twenty-four time points over three growing seasons. Transcriptional and metabolic changes in the kernel elucidate molecular pathways governing nutritional quality, such as the accumulation of unsaturated fatty acids, which are vital for shelf-life and dietary value. This work yields new knowledge and resources that will inform other woody crops and facilitate further improvement of pistachio as a globally significant, sustainable, and nutritious crop.

## Introduction

Tree nuts are the most carbon-efficient protein source of any food^1^. Pistachios are also rich in unsaturated fatty acids, antioxidants, and vitamins^2–7^. Given that pistachio trees are highly resilient to abiotic stress, particularly drought and salinity, they are projected to be an important source of sustainable nutrition in the face of climate change over the next century^8^, with global production of pistachios having more than doubled over the past two decades (FAO; https://www.fao.org/faostat/en/#search/pistachio; **Fig. 1a**).

**Figure 1.**
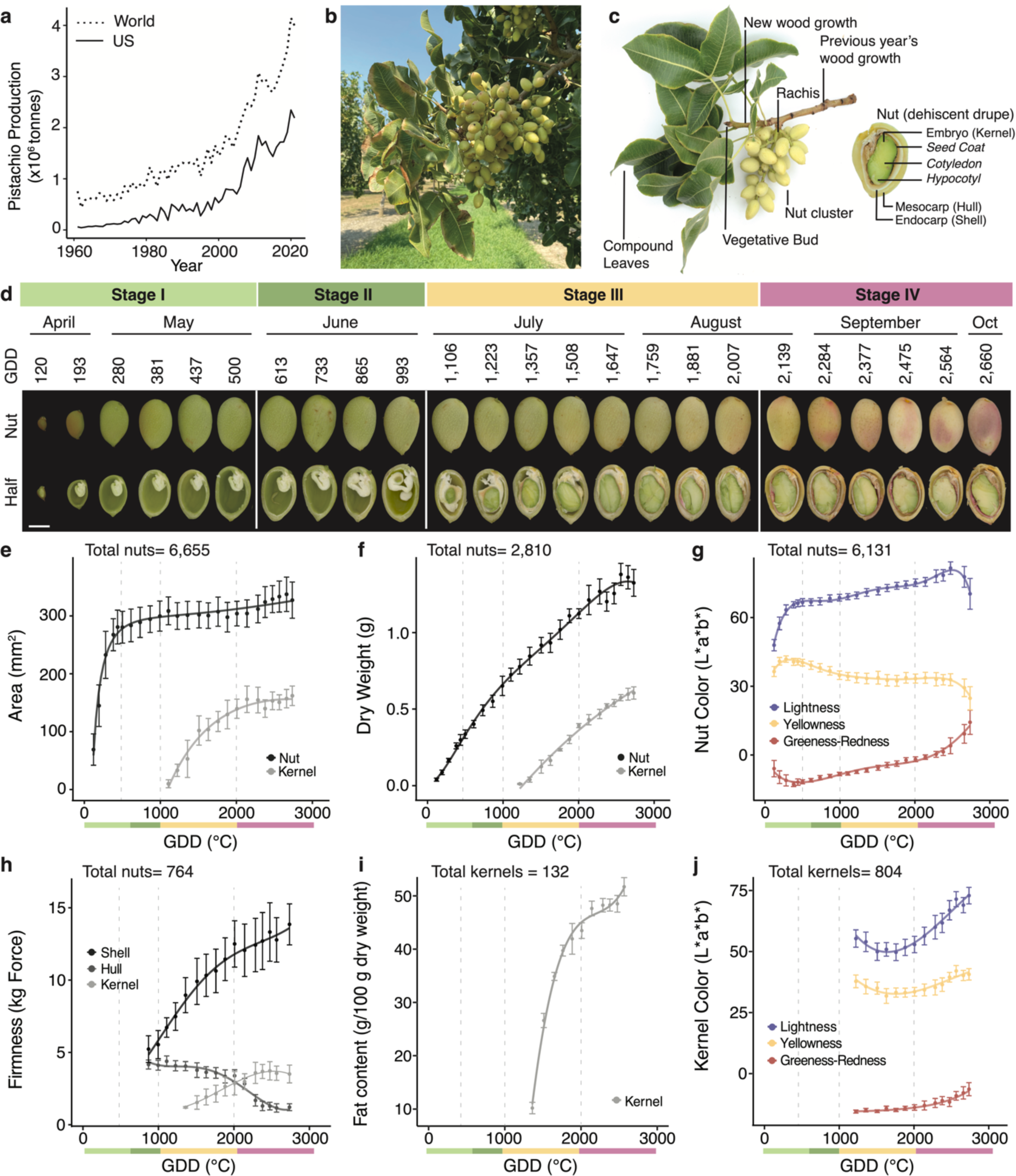
Pistachio nut development is categorized into four distinct stages. **a,** Comparison of United States of America (US) pistachio production to the world in the past 60 years (FAO; https://www.fao.org/faostat/en/#search/pistachio). Pistachios at Stage III on a tree, **b**, and a branch, **c**, with nut and kernel anatomy. **d**, Pistachio development (whole nut, halved nut, and kernel) assessed from April through September 2019 in California and categorized into four stages represented by calendar time and accumulated heat units expressed as growing degree days (GDD) in °C. The new stages were defined by assessing, **e**, whole nut and kernel area growth, **f**, dry weight (g) of the whole nut and kernel, **g**, color changes in the hull measured in the L*a*b* color space, (L*, or lightness, a* or redness, b* or yellowness), **h**, texture changes in the hull, shell, and kernel (kg of Force), (**i**) fat content in the kernel (g/100 g dry weight), and **j**, kernel color changes measured in the L*a*b* color space. **e-j,** Lines show fitted linear and linear-mixed polynomial models as a function of heat accumulation (GDD). **e-j,** The stages are represented in a bar with distinct colors below the x-axes. Stage I=light green, Stage II=green, Stage III=yellow, and Stage IV=pink

Pistachio (*Pistacia vera,* 2n=30) belongs to the Anacardiaceae family, along with cashew and mango, and is the only edible and commercially used species in the genus *Pistacia*. Pistachio trees are dioecious and wind-pollinated. Although commonly known as nuts, pistachio fruits are botanically dehiscent drupes consisting of three main tissues: a leathery exo-mesocarp (hull), a stony endocarp (shell), and an edible seed (kernel) **(Fig. 1b-c)**. Pistachio nut growth has been previously divided into three stages: (1) growth of the hull and shell, (2) shell lignification, and (3) kernel growth^9–13^. These stages, defined by biometric parameters like nut and kernel size, have guided research and agronomic management in the past; yet a complete physiological assessment of pistachio development from fruitset to harvest is lacking.

Also, a better understanding of the molecular mechanisms behind the composition of pistachio kernels can provide a robust foundation for breeding novel pistachio varieties with higher nutritional value and advancing management strategies to boost yield, cut costs, and enhance quality. Such insights provide transferable knowledge that will benefit the development of other hard-shelled nut crops like almonds and walnuts.

Here, we present foundational genomic resources and research into nut development that are critically needed to support the rising demand for pistachio production. We have generated the first chromosome-scale reference-quality genome and annotation of *P*. *vera* ‘Kerman’, the most important female cultivar in the United States, leveraging this resource (available at https://pistachiomics.sf.ucdavis.edu/) to address outstanding questions about the molecular genomics of pistachio nut development. We combined transcriptomic and physiological data to uncover pathways and regulators behind the kernel’s protein accumulation and high unsaturated fat content. Altogether, this work yields a new model of nut development to significantly expand the knowledge of hard-shelled fruit biology, link key molecular processes with the nutritional profile of pistachios, and support growers in decisions like harvest timing and irrigation.

## Results

### Pistachio nuts develop in four distinct stages

Pistachio kernels develop asynchronously from the maternal tissues (hull and shell)^12^ (**Fig. 1c**). To investigate the dynamics of kernel growth in relation to the whole nut, we conducted the most comprehensive study of nut development to date, evaluating physiological traits, such as size and firmness, in pistachio nuts (‘Kerman,’ n=663-6,805) for 24 weeks from fruitset to two weeks after harvest maturity (**Methods**, **Supplementary Table 1, Fig. 1d**). The traits were validated across two additional independent field seasons in distinct geographical locations (**Methods, Supplementary Table 1, Supplementary** Fig. 1**, Supplementary Table 2**). We used both growing degree days (GDD, °C) and calendar-determined time in our analyses because heat accumulation modulates phenology and enzymatic activities that influence metabolism and nut growth^14,15^. These data revealed that four stages define nut development: Stage I (from late April through May in California) occurs when the shell and hull tissues grow in a logarithmic growth pattern plateauing at 500 GDD (**Fig. 1e-f**). During this time, the hull and shell tissues are fused together and display an increasingly green color (**Fig. 1g**). Stage II (June) is the transition period before the beginning of kernel (embryo) growth, when hull and shell expansion stops but the tissues continue to accumulate dry weight (**Fig. 1f**). Stage III (late June to mid-August) corresponds to the kernel growth phase, starting at 1,000 GDD and reaching its maximum size at 2,000 GDD. In contrast to previous reports^9^, we found that shell hardening coincides with kernel growth at Stage III and continues through Stage IV (**Fig. 1e,h**). Stage IV (from late August to September) marks the onset of kernel maturity, at which the kernels reach their maximum size and fat content despite the continued increase in dry weight (**Fig. 1i**). Kernel maturity coincides with hull ripening (*e.g.,* tissue softening and color changes) and shell split or dehiscence. At this stage, the kernels start losing their deep green coloration, preparing for seed dormancy (**Fig. 1g,h**). These findings show that pistachio nut development is closely regulated by heat accumulation, enabling accurate prediction of growth stages across seasons.

### A reference quality genome and annotation of pistachio

Our study aimed to characterize the molecular genomics of pistachio nut development. To enable this, and future pistachio research more generally, we generated an ultra-high quality chromosome-scale genome assembly and annotation for *Pistacia vera* ‘Kerman’. This reference genome for pistachio surpasses previous efforts in accuracy and completeness, providing a foundation for breeding and -omics research **(Table 1)**. Our assembly illuminates previously undescribed details of the pistachio genome architecture, particularly the presence of repetitive, knob-like regions on eleven chromosomes **(Fig. 2c)**. These regions, characterized by megabase-scale 178 bp satellite repeats, reveal a remarkable genomic feature that had been observed cytologically^16^, but was previously obscured from genome assemblies. This discovery reshapes our understanding of the pistachio genomic landscape and highlights the value of robust genome assembly for molecular genomic and evolutionary research.

**Figure 2.**
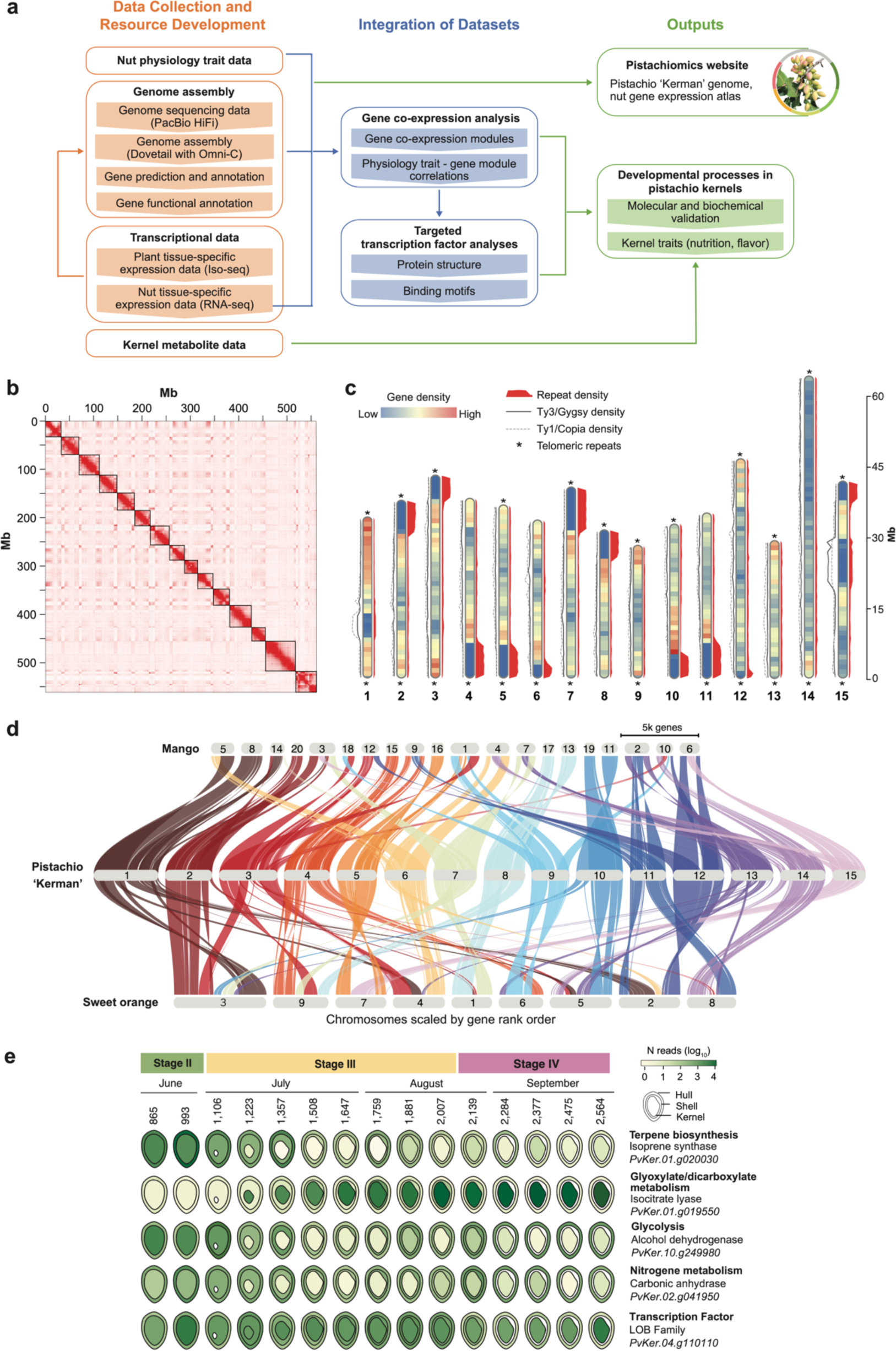
Chromosome-scale genome assembly of *Pistacia vera* ‘Kerman’ offers new genetic resources and tools. **a**, Overview of current study workflow from data collection to outputs. **b**, Heat map of the Omni-C interaction density among 15 chromosomes. The red color indicates the intensity of interactions between genomic regions. **c**, Ideogram with protein-coding gene and repeat density in 1 Mb window size on 15 chromosomes of the ‘Kerman’ genome. Protein coding gene density is represented in each chromosome in heatmap style, and repeat density is plotted to the right side of each chromosome in red. The density of long terminal repeat retrotransposons (LTR-RTs) Ty3/Gypsy and Ty1/copia are shown on the left side of each chromosome in normal and dotted lines, respectively. The scale bar for chromosome size is indicated on the right in light gray. **d**, Macrosynteny analysis of the ‘Kerman’ 15 chromosomes compared to the mango and sweet orange genomes. **e**, Tissue-specific RNA-seq expression of genes highly expressed unique to pistachio nuts identified by comparison of Iso-seq collapsed isoforms demonstrating differential expression patterns across tissues.

**Table 1.** Characteristics of the *Pistacia vera* ‘Kerman’ genome and annotation.

| *P. vera* 'Kerman' |  |  |
| --- | --- | --- |
| Genome assembly | Primary assembly | Chromosomes |
| Number of Seq. | 102 | 15 |
| Size (bp) | 579,837,297 | 559,112,681 |
| Max length (bp) | 49,785,438 | 62,545,632 |
| N50 (bp) | 25,977,840 | 36,813,375 |
| L50 | 9 | 7 |
| L90 | 21 | 13 |
| Number of Gaps |  | 12 |
| %GC | 37 | 36.67 |
| BUSCO completeness (%) | 98.8 | 98.9 |
| complete single-copy (%) | 95.4 | 95.8 |
| complete duplicated (%) | 3.3 | 2.8 |
| Protein-coding gene | Chromosomes |  |
| # of genes | 37,063 |  |
| Total length of genes (bp) | 108,162,188 |  |
| % of genome covered by genes | 19.35 |  |
| Mean gene length (bp) | 2,918 |  |
| Mean CDS length (bp) | 1,073 |  |
| Mean exon length (bp) | 285 |  |
| Mean intron length (bp) | 436 |  |
| Mean exon number per gene | 4.6 |  |
| BUSCO completeness (%) | 98.9 |  |
| complete single-copy (%) | 97.2 |  |
| complete duplicated (%) | 1.7 |  |
| Non-coding loci | Number | Total (bp) |
| tRNA | 514 | 38,570 |
| rRNA | 848 | 895,039 |
| miRNA | 182 | 24,122 |
| snoRNA | 2,123 | 223,030 |

We resolved the ‘Kerman’ genome to 15 chromosomes (559.11 Mb) with only 12 gaps, by scaffolding and manually curating 102 non-organelle contigs (579.8 Mb) with Omni-C data **(Fig. 2b,c and Table 1, Supplementary Table 4)**. The k-mer analysis with PacBio HiFi reads **(Supplementary Table 3)** estimated the ‘Kerman’ genome size to be approximately 521 Mb, with moderate heterozygosity (0.755%) and repetitiveness (54.1%) **(Supplementary** Fig. 2). Notably, our assembly has a significant reduction in the number of duplicated complete BUSCO genes compared to previous reports, from between 210-286 (previous) to only 45 (this study), indicating the elimination of false duplications **(Supplementary Table 5)**.

The ‘Kerman’ genome contains over 0.83 million repetitive elements constituting 376.56 Mb (∼65%) of the assembled genome **(Supplementary** Fig. 3 **and Table 6)**. This notable increase from previous genomes (53% and 56%) **(Supplementary Table 6)** implies accurate recovery of repetitive regions while successfully excluding false-segmental duplications explained by smaller assembly size (559.11 Mb) than the other assemblies, ‘Batoury’ (671 Mb)^17^, and ‘Siirt’ (596 Mb) and ‘Bagyolu’ (623.4 Mb)^18^. Consistent with other plant species^19,20^, long terminal repeat retrotransposons (LTR-RTs) were the most abundant (48.95%) class in the ‘Kerman’ genome. We also discovered regions containing massive enrichments of 178 bp repeats known as PIVE-180 on one arm of eleven chromosomes in pistachio (*e.g.,* up to ∼9 Mb on chromosome 7, where no protein-coding genes were present) **(Fig. 2c)**. Importantly, the presence of these regions has been confirmed by previous cytological work, with the location of these repetitive sequences being distinct from centromeres and corresponding with those observed in our genome assembly^16^. The existence of these extremely repetitive satellite DNAs likely limited the accuracy of previous genome assemblies and chromosome construction. The syntenic analysis showed overall collinearity but a noticeable presence and absence of variation of these and other genomic regions, which implies a significant improvement in the quality of the pistachio genome **(Supplementary** Fig. 4).

We annotated 37,955 protein-coding genes in the ‘Kerman’ genome with multiple lines of evidence (**Materials and Methods)**. Among these, multiple isoforms were characterized in 6,938 genes, and 22,942 genes were functionally annotated with multiple databases (**Materials and Methods**, **Supplementary Table 4)**. The majority of genes (>97%, 37,063) were anchored onto the 15 chromosomes **(Table 1)**. The quality of gene annotations reached a near 99% completeness in BUSCO value **(Supplementary Table 8)**. A complete gene annotation, essential for our study of pistachio nut development and other applied pistachio research, was achieved with high-quality assembly, long and short-read transcript evidence from various tissue types, and *ab initio* predictions **(Supplementary** Fig. 5 **and Table 8, Materials and Methods)**. The genome assembly and annotation of *P*. *vera* ‘Kerman’ enabled more accurate orthology-constrained macrosynteny analysis between genomes of pistachio and the close relatives mango and sweet orange, revealing extensive structural variation even within the family and whole genome duplication in the mango genome **(Fig. 2d)**.

### Spatiotemporal gene expression data support the distinct nut developmental stages

The high-quality genome assembly and annotation of ‘Kerman’ provided a resource to investigate nut development at the molecular genomic level. We measured tissue-specific whole transcriptome gene expression across 15 weeks of nut development encompassing kernel growth and maturation (the end of Stage II through Stage IV; **Fig. 1d**, **Fig. 2e**). These data are available as an atlas of spatiotemporal expression (https://pistachiomics.sf.ucdavis.edu/).

The kernels used in this study resulted from open pollination of ‘Kerman’ (maternal) with Peters (paternal; the male cultivar used as a pollenizer in ‘Kerman’ orchards). Based on their transcriptional patterns, we distinguished the maternal tissues (hull and shell) from the hybrid kernels, reflecting differences in genetic backgrounds and ontogenic states (**Fig. 3a**, PC1). We also observed differences in transcriptional patterns across nut development stages (**Fig. 3a**, PC2).

**Figure 3.**
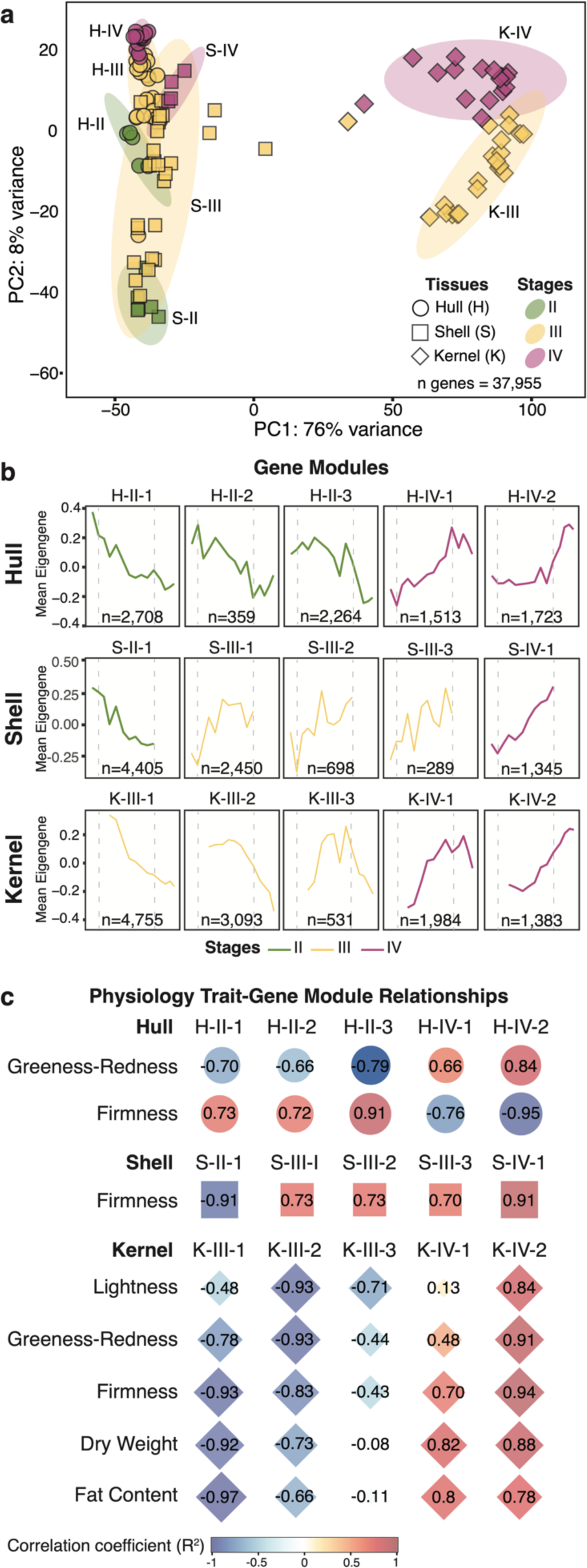
Gene co-expression patterns confirm pistachio developmental stages. **a,** Principal component analysis (PCA) of total gene expression (normalized reads) for all samples, marked by stage (color) and tissue (shape). Then, a weighted gene co-expression network analysis was conducted for each tissue and produced modules of genes with similar expression patterns. **b,** Gene modules were selected with high correlations to physiological traits and categorized by stage according to the time points in which expression was elevated (mean eigengene value) in each tissue type (H-hull, S-shell, or K-kernel) and at specific developmental stages (II, III, or IV) for each module (1-X). Gray dashed lines indicate the transitions between stages. All gene modules can be found in Supplementary Table 10. **c,** Correlation between the expression profiles of the selected modules and the physiological trait data. Correlations R^2^ >0.7 with significance *P*<0.01 are shown for each tissue. The intensity of the color indicates the strength of the correlation (R^2^), and the shape of the icon indicates the tissue type.

We identified groups of genes, *i.e.*, co-expressed gene modules, in each tissue (H-hull, S-shell, K-kernel) with high expression at specific developmental stages (I-IV) that strongly correlated with the occurrence of physiological changes, providing critical insights into the link between molecular processes and the four nut developmental stages (**Fig. 3b-c, Supplementary** Fig. 6**: Modules, Supplementary Table 9**). Fat content in the kernel was highly correlated to K-III-1, which was enriched for ‘fatty acid biosynthesis’ and ‘fatty acid elongation’ functions. Shell lignin biosynthesis genes were highly expressed and enriched (‘phenylpropanoid biosynthesis’) in the S-II-1 gene module, correlating to early shell hardening (Stage II-III) when lignin is deposited (**Supplementary Table 10**). Similarly, the cell wall degrading enzymes pectate lyase and a β-glucosidase potentially involved in hull softening were the top expressed genes in their respective modules, H-IV-1 and H-IV-2, coinciding with hull ripening in Stage IV (**Fig 3c**).

### Gene modules reveal pathways involved in kernel nutritional quality

The molecular events occurring in the kernel during Stages III and IV led to the gain of nutritional traits (**Fig. 4a**). During Stage III, kernels had a deep green color and grew rapidly (**Fig. 1f,j**). We identified the gene module K-III-1 to be highly correlated with color and the rapid growth during Stage III and found enrichments in functions involved in energy metabolism (‘photosynthesis’), carbohydrate metabolism (‘starch and sucrose metabolism’), and protein accumulation (‘amino acid metabolism’) (**Supplementary Table 10**). This was supported by increased protein, carbohydrate, and starch content in the kernels during Stage III, with protein making up the highest proportion of these macronutrients (**Fig. 4a,b**). The enrichment of fatty acid degradation genes, specifically of ɑ-linoleic acid, coincided with the plateau of fatty acid content measured during Stage IV. Further, the ratio of mono-unsaturated fatty acids (MUFA) to poly-unsaturated fatty acids (PUFA) increased during Stages III and IV (**Fig. 4a,c**).

**Figure 4.**
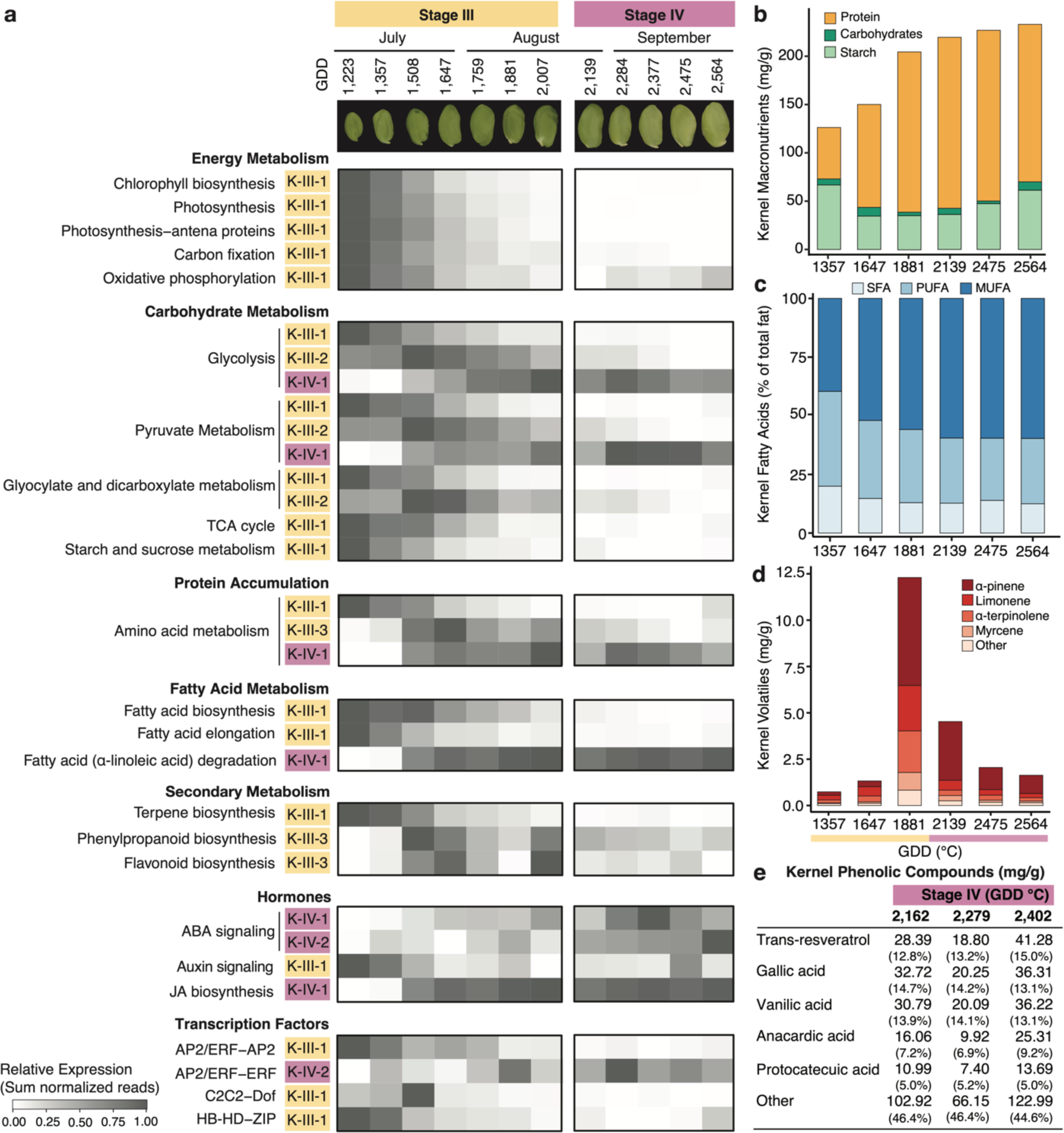
Pistachio kernels display conserved patterns of seed development and unique metabolite fluctuations. **a,** Summary of significantly enriched gene functions (*e.g.,* GO, KEGG, iTAK) in modules highly correlated to relevant kernel traits (Supplementary Table 10: Enrichments). The sum of gene expression (*i.e.,* normalized reads) at each time point in a given module is shown for enriched (*Padj*<0.05) functions. Metabolite profiles to confirm the gene expression trends across kernel development were obtained for **b**, carbohydrates, starch, and proteins, **c**, fatty acids including mono-unsaturated fatty acid (MUFA), polyunsaturated fatty acid (PUFA), and saturated fatty acid (SFA), **d**, monoterpene volatiles relevant to flavor, **e**, phenolic compounds contributing to flavor and nutrition. Phenolics are reported as mg/g dry weight and as a percentage of the total amount. Metabolites are reported as averages across kernel samples (n=3-4). The time points of kernel development correspond to the growing degree days (GDD, °C) at collection.

Production of volatile terpenoids and flavonoids in the kernel produces flavor and antioxidant compounds, respectively. We observed a large peak in monoterpenes at the start of Stage IV (1881 GDD), shortly after peak expression of terpene biosynthesis genes (**Fig. 4a, d, Supplementary Table 11**, **Supplementary Table 9**). The rate-limiting enzyme geranyl-diphosphate synthase (GPS) and monoterpene synthases were among the genes co-expressed during Stage III (primary K-III-I), which can explain the large increase in volatiles in the transition to Stage IV (**Supplementary Table 9**). At harvest, ɑ-pinene was the main monoterpene present in the kernels, consistent with previous findings^5,7^. Total phenolic compounds increased during Stage IV, supporting claims and previous reports that pistachio kernels can be an important source of antioxidants^5,7^. Likewise, phenylpropanoid biosynthesis (K-III-3 and K-IV-6) and flavonoid biosynthesis (K-III-3) were enriched and genes encoding the rate-limiting steps phenylalanine ammonia-lyase (PAL) and chalcone synthase (CHS) were expressed in each respective module. Unlike studies in other varieties, trans-resveratrol (41.28 mg/g) was the most abundant among measured phenols, indicating it may be a major antioxidant in ‘Kerman’ kernels^21–23^ (**Fig. 4e**, **Supplementary Table 11)**.

### Conserved transcriptional regulators are associated with fatty acid accumulation in pistachio kernels

Abscisic acid (ABA) signaling components, including homologs of PYR, SnRK2, ABF, and PP2C, were enriched and highly expressed in modules K-IV-1 and K-IV-2 (**Fig. 4a**). ABA is a key regulator of seed maturation leading to seed dormancy^24^. We validated that the concentration of ABA was higher in the kernel (2,375.578 mg/g) relative to the maternal tissues (681.019 mg/g in the hull and 56.474 mg/g in the shell) at 1,881 GDD when the nut is transitioning to Stage IV (**Supplementary Table 12**). Jasmonic acid (JA) biosynthesis, derived from fatty acid ɑ-linoleic acid, was also enriched with increased expression during Stage III and IV (**Fig. 4a**). Homologs of the JA signaling transcription factors (TFs) JAZ and MYC2 were highly expressed in module K-IV-1. Although we detected JA in the kernels (6 mg/g at 1,881 GDD corresponding to late Stage III), the hull and the shell had higher levels of this hormone at the same stage (1,700 mg/g and 33 mg/g, respectively).

We mined TFs with high expression (top 10%) among the gene modules to identify potential regulators of kernel development. In particular, the module K-III-1 enriched in fatty acid biosynthesis and elongation (**Fig. 4a**) included the homolog of one of the four major seed master TFs, *LEAFY-COTYLEDON1* (*NFYB-LEC1*), known to co-regulate fatty acid biosynthesis across diverse crop species^25^. This module also contained the homolog of a key regulator of seed oil content, *WRINKLED-1* (*AP2-WRI1*)^26^, which has been reported to be under the control of NFYB-LEC1. The *PvAP2-WRI1* gene in the ‘Kerman’ genome had a high degree of protein sequence similarity and conservation of DNA-binding domains across 16 representative angiosperm taxa^27–29^ **(Fig. 5b, Supplementary** Figs. 7**, 8)**. Fatty acid biosynthesis genes in the K-III-1 module had the putative AW-box binding domain significantly closer to the translational initiation site compared to other genes in the genome that contained the same sequence, suggesting that PvAP2-WRI1 may regulate them, but this result will require experimental validation^30^ **(Fig. 5c)**. The AW-box and the conserved binding motif (CCAAT) for the PvNFYB-LEC1 predicted protein were prevalent among genes involved in *de novo* fatty acid biosynthesis steps and desaturase enzymes associated with the synthesis of MUFA and PUFA (**Fig. 5d)**.

**Figure 5.**
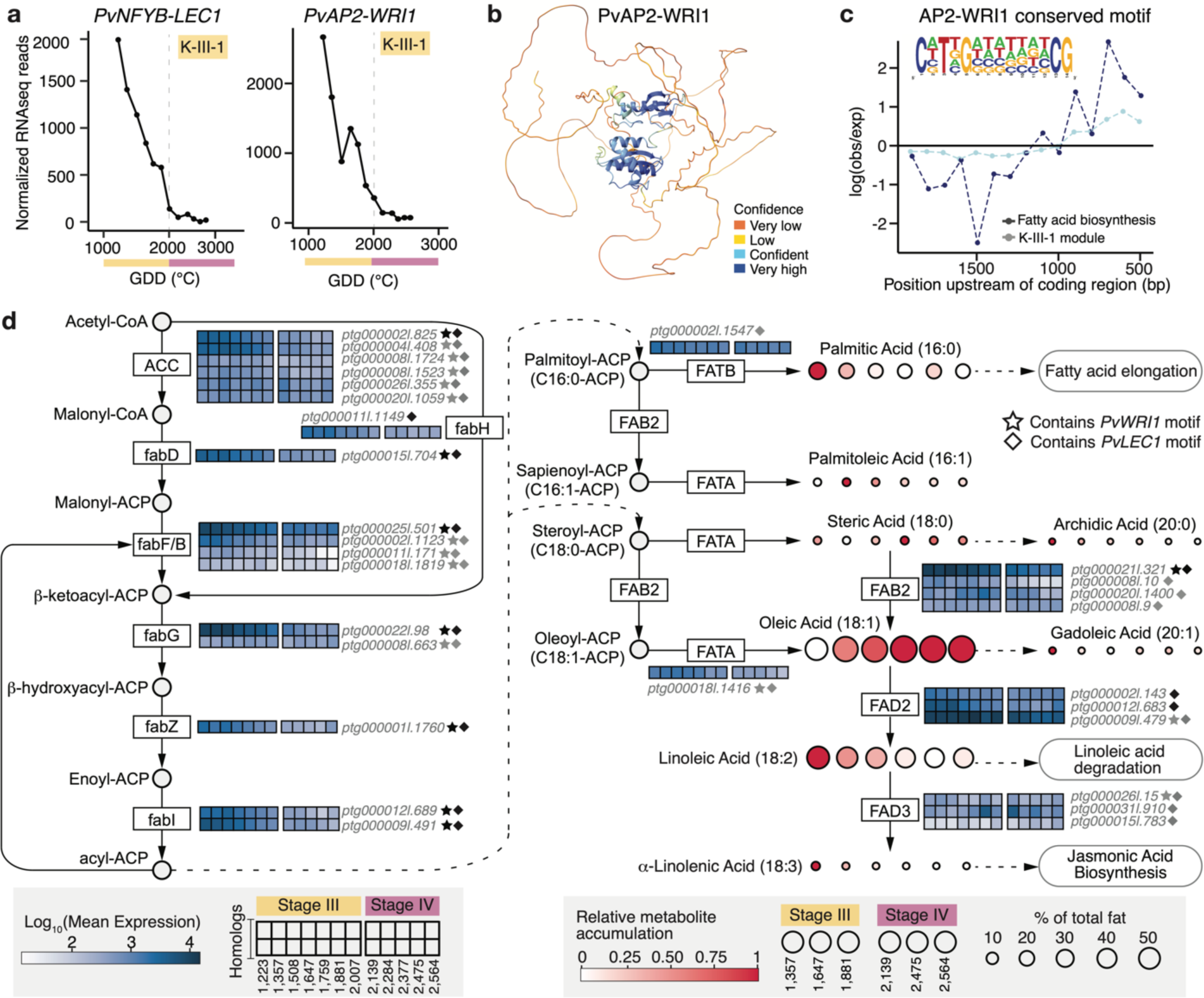
Fatty acid biosynthesis and potential regulators in pistachio kernel development. **a,** Expression pattern of transcription factors *LEAFY-COTYLEDON1* (*PvNFYB-LEC1*; *PvKer.12.g280410*) and *WRINKLED1* (*PvAP2-WRI1*; *PvKer.03.g083330*) involved in seed development and fatty acid accumulation. **b**, Predicted 3D structure of *P*. *vera* ‘Kerman’ PvAP2-WRI1 protein. Colors indicate the folding confidence level as labeled on the right bottom. **c**, Relative location of previously reported WRI1 AW-Box binding motifs found in ‘Kerman’ fatty acid biosynthesis (dark blue) and K-III-1 module genes (light blue) compared to other genes that contained the same binding motif. The consensus sequence of the AW-Box binding motif is shown as a logo. **d,** A representation of fatty acid biosynthesis pathways based on KEGG pathways (www.genome.jp/kegg/pathway.html, last accessed December 2023). Dashed lines indicate that some steps were omitted. The gene expression levels (*i.e.*, log10 of the mean normalized read count for each sampling) are represented in colored boxes, with each box representing a sampling point from Stage III and Stage IV determined by growing degree days (GDD, °C) and each row representing a gene. Samplings for gene expression include dates from 1,223 GDD to 2,564 GDD. Colored circles represent the accumulation of specific fatty acid compounds. Colors indicate the changes in metabolites through time, and the size of each circle represents the amount of the metabolite, measured in normalized area percentages (%). Three time points from Stage III (1357, 1647, 1881 GDD) and Stage IV (2,139, 2,475, 2,564 GDD) were used for all fatty acid data shown. Fatty acid genes containing the WRI1 AW-box or LEC1 CCAAT target sequences are indicated by a star or diamond shape, respectively. ACA, acetyl-CoA carboxylase; fabD, S-malonyltransferase; fabF, 3-oxoacyl-[acyl-carrier-protein] synthase II; fabH, 3-oxoacyl-[acyl-carrier-protein] synthase III; fabG, 3-oxoacyl-[acyl-carrier protein] reductase; fabZ, 3-hydroxyacyl-[acyl-carrier-protein] dehydratase; fabI, enoyl-[acyl-carrier protein] reductase I; FATB, fatty acyl-ACP thioesterase B; FATA, fatty acyl-ACP thioesterase A; FAB2, acyl-[acyl-carrier-protein] desaturase; FAD2, omega-6 fatty acid desaturase; FAD3, acyl-lipid omega-3 desaturase.

To further understand the dynamics of fat accumulation in pistachio, we visualized the expression of fatty acid biosynthesis genes across kernel development and measured individual metabolites (**Fig. 5d**). The highest expressed genes in the pathway were the *FAB2* homolog, an acyl-[acyl-carrier-protein] desaturase, and the FAD2 homolog, an omega-6 fatty acid desaturase. These genes represent key steps in unsaturated fatty acid biosynthesis and contain the WRI1 and LEC1 binding motifs^25,31,32^. The metabolite data showed that oleic acid, a MUFA, and linoleic acid, a PUFA, were the top two accumulated fatty acids in pistachio kernels. The relative accumulation of linoleic acid decreased over time while oleic acid increased, as was previously observed^5^, and may be due to fatty acid degradation happening in late kernel development (K-III-3 and K-IV-1; **Fig. 4a**).

## Discussion

This comprehensive study positions pistachio to serve as a model system to investigate the biology of hard-shelled fruits and the asynchronous behavior of seed and fruit development observed in some tree species^32^. The ‘Kerman’ genome produced here provides a foundational resource for pistachio breeding and sets the stage for further molecular studies and deeper investigations into the notable genomic features discovered, particularly large knob-like satellite repeats found on multiple chromosomes. Leveraging this genome, we were able to map molecular processes underlying nut development at high spatiotemporal resolution.

Pistachio kernels do not grow until approximately 1,000 GDD after maternal tissue growth starts (**Fig. 1d-j**). Energy limitations in the tree may explain why this pattern has evolved in some tree species. Carbohydrates reserved from the previous year are exhausted after developing the hull, shell, and leaves in Stage I^33^. The lull in fruit growth identified as Stage II may function to generate resources from photosynthesis that can be used to support kernel growth in Stage III^34^. This growth pattern may also help explain why some pistachio nuts continue their development without a kernel (*i.e.,* “blanks”) and why the trees drop their buds in early summer (*i.e.,* bud abscission), leading to alternate bearing years^35,36^. Transcriptome and hormone analysis at the transition to seed growth (Stage II to III) will be essential to further elucidate these phenomena.

Transcriptomic analysis during kernel growth and maturation highlighted that pistachio has conserved regulatory mechanisms during seed development consistent with other plant species (**Fig. 4**). Paralogs of the master transcription factors critical to seed development known as LAFL (LEC1, ABI2, FUS3, and LEC2) were highly expressed and followed conserved expression patterns during pistachio kernel development (**Supplementary Table 9)**. Further, we identified homologs of LEC1 and WRI1, known to act together to regulate seed oil content in other plant species^25,31^, to be candidate regulators of fatty acid biosynthesis in pistachio^37^. Benny and colleagues^35,36^ found *WRI1* and coregulators *SUCROSE-NON-FERMENTING1-RELATED PROTEIN KINASE1* (*SnRK1),* and *trehalose 6-phosphate (T6P)* were co-expressed in early kernel development of ‘Bianca’ pistachios. Likewise, *WRI1*, *T6P* and *SnRK1* were in the K-III-1 module associated with increased fat content and should be further explored to define the regulatory network involved.

The knowledge and genomic resources provided here will facilitate molecular-assisted breeding and the study of the biological causes of pistachio quality defects, like internal kernel discoloration (IKD), low shell split, and hull deterioration. Further, our research can be readily applied to pistachio production. The newly defined Stage II, a crucial period of reduced nut growth before kernel initiation, has significant implications for management practices such as the timing of deficit irrigation^9^. We discovered that shell hardening and kernel growth occur simultaneously, providing a non-destructive benchmark to track kernel development, corroborating an earlier report by Zhang and colleagues^10^. The hull ripening traits observed during Stage IV can give non-destructive indications of the best harvest time. Kernels reach their maximum ratios of macronutrients at the start of Stage IV, while changes to volatiles and phenolics occur later; this should be considered to achieve the best flavor and nutrient content at harvest. Overall, as the demand for pistachios continues to increase, our work provides multiple avenues for research that can benefit breeders, growers, and consumers.

## Methods

### Sample collection for physiological data, RNAseq experiments, and metabolic analysis

Pistachio physiological data were collected across three growing seasons (2019, 2020, 2021) for evaluation from an experimental pistachio orchard (cv. ‘Kerman’ grafted onto UCB1 rootstock) with 30-year-old trees at the University of California Kearney Agricultural Research and Extension Center (Fresno County) in 2019. The results were validated in 2020 in a commercial orchard with 10-year-old trees of the same cultivar located in Woodland, CA (Yolo County) and again in 2021 in a separate commercial orchard with 10-year-old trees located in Three Rocks, CA (Fresno County). In all the orchards the pollinizer cultivar was ‘Peters’. In each study, homogenous trees (n_2019_=12, n_2020_=4, n_2021_=8) were randomly selected across the orchard and were continuously sampled throughout the season. In 2019, samplings were conducted weekly after fruit set on April 25th (120 growing degree days (GDD) (°C), 10 days post anthesis (dpa) continuing through October 14th (2760 GDD), corresponding to two weeks after harvest maturity (70% shell split). About 48 clusters were collected per time point from 12 trees. In 2020, samplings occurred weekly from April 19th (24 GDD) through September 10th (2452 GDD, commercial harvest), and in 2021, samplings occurred weekly from July 15 (1531 GDD) to September 22 (2828 GDD). Four whole clusters yielding about 50 nuts of uniform maturity were collected per tree at each sampling. Environmental temperature was recorded using HOBO temperature and light sensors data logger (Onset, United States) placed in the orchards at three separate locations. Growing degree days were calculated with the equation^10^:

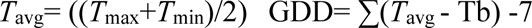

Samples were dissected into each tissue (hull, shell, and kernel) and frozen in liquid nitrogen on the day of sampling. The frozen tissues were ground into a fine powder with the Retsch Mixer Mill MM 400 (Verder Scientific, Netherlands).

### Physiological measurements and statistical modeling

To assess nut area and color, nuts were imaged longitudinally using a VideometerLab 3 (Videometer, Denmark) facilitated by Aginnovation LLC. VideometerLab 3 software was utilized for image analysis in both 2019 and 2020. Color measurements were taken on the L*a*b* color scale as an average across the entire nut area. For color and area, 10-30 nuts per tree (n=12) were sampled weekly for 25 weeks. Growth was determined through fresh weight and dry weight measurements of the whole nut and the kernels. Fresh weight was taken on the day of harvest of 10 nuts per tree (n=12), and the average per nut weight was calculated. Nuts were cut open and divided into kernels and the remaining tissues (hull and shell). Nuts were placed in a drying oven at 80℃ for 2 days until all moisture was evaporated and measured. The average per nut and per kernel weights per cluster (n= 7-12 per sampling) were calculated from the total weight and total number of nuts. Shell split was measured as the incidence of nuts with any degree of separation between sides and taken as a proportion of the total nuts. Destructive texture measurements were obtained with the use of a TA.XT2i Texture Analyzer (Texture Technologies, United States) using a TA52 2 mm probe with a trigger force of 5 g and a test speed of 2.00 mm/sec with Exponent software (Texture Technologies Corporation, United States). The probe punctured through the hull, shell, and kernel tissues, and the peaks of each tissue were distinguished and recorded in the software. Measurements were reported as kilograms (kg) of force. 20 to 60 nuts were assessed every sampling for 25 weeks. Fat content was obtained from oven-dried kernels, as described by Polari and colleges^38^. Briefly, dried pistachio kernels (5.0 ±0.1 g) were ground in a coffee grinder (JavaPresse Coffee Company, United States), weighed into a cellulose extraction thimble, placed in the Soxhlet extractor, and extracted using n-hexane for 6 h. The solvent was distilled, and the residual solvent was eliminated in an oven at 105 ℃ for 3 h. Fat content was expressed as grams of fat per 100 grams.

Physiological traits, including kernel and nut dry weights (g), kernel and nut areas (mm^2^), kernel and nut colors (L*, a*, b*), and kernel, shell, and nut textures were modeled against heat accumulation (growing degree days, °C). Various Box-Cox transformations (i.e. square, square root, and logarithm transformation, etc.) on traits data using the MASS package ^39^ in R were made before modeling to ensure an approximate normal distribution of traits and roughly equal variance of the error terms. Linear and linear mixed models were fitted for each trait with a polynomial of 2 or 3 as a function of heat accumulation, using lme4 and lmerTest packages in R^40,41^. Random intercepts were added in linear mixed models. Random effects included cluster, tree, and year, depending on the models **(Supplementary Table 2: Models)**.

### Sample collection, nucleic acid extraction, library construction, and sequencing for PacBio HiFi, Iso-Seq, and Omni-C sequencing

High molecular weight genomic DNA was extracted from young leaves of ‘Kerman’ trees located at Finca “La Entresierra,” Centro de Investigación Agroambiental “El Chaparrillo” (CIAG), Spain, using Circulomics Nanobind Plant nuclei kit (http://circulomics.com, Pacific Biosciences, United States) after nuclei isolation according to Workman and colleagues^42^. The library construction and PacBio HiFi sequencing were completed on a Sequel II system at Gentyane facilities in the French National Institute of Agronomy (INRAE). As a result, a total of 25.18 Gb PacBio HiFi reads with a mean length of 16.3 kb was generated **(Supplementary Table 3)**. For Omni-C data, young leaf tissue samples were collected, directly snap-frozen in liquid nitrogen, stored at −80℃, and sent to Dovetail genomics for the Omni-C library construction and sequencing.

For Iso-seq sequencing, samples were collected from the same ‘Kerman’ tree used for genome assembly at specific developmental stages and immediately frozen in liquid nitrogen. The collection dates were as follows: dormant buds on March 23, breaking buds and flowers on April 6, leaves on April 16, and fruits on May 6, 2021. 100 mg of material for each sample was pulverized with a pestle and mortar in liquid nitrogen. Total RNA was extracted by using the Spectrum Plant Total RNA (Sigma-Aldrich, United States) Kit, according to the protocol recommended for “difficult” species. The quantity and purity of the total RNA were checked with the Nanodrop (ND1000 Thermo Fisher Scientific, United States). Iso-seq library prep and sequencing were performed using RNA samples with ∼ 400 ng/microliter for the different tissues that were barcoded and sequenced in the same Sequel II SMRT cell at Gentyane facilities in INRAE.

### Genome size estimation

Genome size, heterozygosity, and repeat content were estimated based on k-mer frequency analysis with PacBio HiFi reads. Jellyfish v2.2.10 41 ^43^ was used to count 21-mers with a maximum *k*-mer depth of 1e6, which takes repetitive regions into account. The resulting histogram from Jellyfish was subjected to Genomescope v1 web^44^ to estimate genome size, levels of heterozygosity, and repeat content.

### Genome assembly and chromosome construction

A *de novo* assembly of PacBio HiFi reads into contigs was performed using Hifiasm v0.16.0^45^ with default parameters. Organelle (plastid and mitochondrial) origin contigs were filtered out from the primary contig assembly using ‘Kerman’ plastid and mitochondrial sequences that were pulled out from PacBio HiFi reads in Geneious Prime (https://www.geneious.com). The high-coverage Omni-C data (99.92 Gb and ∼172X depth) was quality-checked with FastQC toolkit^46^ and aligned to filtered primary contig assembly using Juicer v1.6^47^. These aligned read pairs were utilized to scaffold the assembly into 15 chromosomes based on chromatin interaction information using 3D-DNA^48^ with default parameters and manual correction on Juicebox^49^. The filtered primary contig assembly was scaffolded again with 3D-DNA scaffolds as a reference using RagTag v2.1.0 with default parameters^50^ followed by manual correction by comparing both RagTag and 3D-DNA scaffolds. Chromosome numbers and orientations from previously published pistachio genomes were used^18^. The completeness of assemblies, scaffolds, and 15 chromosomes was assessed using BUSCO (Benchmarking Universal Single-Copy Orthologs) v5.4.4^51^ with the embryophyta_db10 database. The overall workflow from sample collection to the chromosome-scale genome is provided in **Supplementary** Fig. 9.

### Genome synteny analysis

The complete genome and annotation of *Mangifera indica* (mango)^52^ in the same family, Anacardiaceae, and *Citrus sinensis* (sweet orange)^53^ in the same order, Sapindales were downloaded to conduct a comparative analysis. Synteny of the chromosome-level genome of ‘Kerman’ was compared with that of Mango and Sweet Orange using GENESPACE v0.9.4^54^, which takes orthology information into account using OrthoFinder v2.5.4^55^. The manual reformation of gene annotation and protein files was needed for GENESPACE input.

### Genome annotation

Annotation of transposable elements (TE) was accomplished with primary contig assembly using Extensive de-novo TE Annotator (EDTA) v2.1.0^56^, which generates a non-redundant TE library and TEs classified into the superfamily level. Sequences from the non-redundant TE library overlapping with the filtered plant protein database were filtered out using protExcluder v1.2 (https://www.canr.msu.edu/hrt/uploads/535/78637/ProtExcluder1.2.tar.gz) to hinder exclusion of genes in the gene prediction analysis. Output from protExcluder was employed to re-annotate TEs in the primary contig assembly using EDTA. The assembly was softmasked for further gene prediction analysis.

To annotate genes, softmasked ‘Kerman’ assembly, RNA sequencing (RNA-seq), and Isoform sequencing (Iso-seq) data were used. PacBio Iso-seq data was assembled from five different tissues using the IsoSeq3 pipeline (Pacific Biosciences, United States) with the following steps: i) demultiplexing and primer removal using lima v2.0.0, ii) removing of poly(A) tails and concatemers, iii) clustering isoforms, and iv) mapping to the assembly using pbmm2 v1.9.0 and collapsing isoforms. The quality of RNA-seq data from pistachio fruit (hull and shell) was assessed using FastQC v0.11.9^46^. Raw RNA-seq reads were filtered and adapter trimmed using Trimmomatic v0.39^57^ with the following parameters: maximum seed mismatches = 2, palindrome clip threshold = 30, simple clip threshold = 10, minimum leading quality = 3, minimum trailing quality = 3, and minimum length = 36, and clean reads were mapped to the assembly using Hisat2 v2.2.1^58^. Iso-seq demultiplexed reads were aligned to the assembly using minimap2^59^. Aligned RNA- and Iso-seq reads were independently assembled into transcripts using StringTie v2.2.1^60^. Before *ab initio* gene prediction, the aligned Iso-seq data and softmasked assembly were used as a retraining set for Braker2 v2.1.2^61^. Augustus v3.1.0^62^ was used to make the *ab initio* prediction with retraining files, softmasked assembly, and exon hints from Iso-seq isoforms produced in the IsoSeq3 pipeline. All data, including assembled RNA- and Iso-seq data and Augustus output gene models, were combined to find consensus gene models using EvidenceModeler (EVM) v2.0.0^63^ with different weights for each input data (7, 4, and 1 for Iso-, RNA-seq transcripts, and Augustus gene models, respectively). Finally, 5’ and 3’ UTR regions and different isoforms were updated from EVM gene models using PASApipeline v2.5.2^63^ based on the alignment of Iso-seq demultiplexed reads. The workflow of the gene annotation pipeline mentioned above is provided in **Supplementary** Fig. 10. The final gene annotation on primary contig assembly was lifted over to 15 chromosomes, and gene IDs were curated using modified python script GFF_RenameThemAll.py^64^. The quality of the final gene models was assessed using BUSCO^51^ (embryophyta_db10 set) with protein sequences extracted by using AGAT (Another GTF/GFF Analysis Toolkit) v0.9.1^65^. The RIdeogram R package^66^ was used to plot gene and TE density on the 15 chromosomes in the ‘Kerman’ genome assembly. The 3D structures of gene models were characterized in an atomic resolution with a language model using ESMFold^67^ and visualized using UCSF ChimeraX v1.6.1^68^.

tRNAs genes were identified using tRNAScan-SE v2.0.12^69^ with eukaryote parameters. rRNA genes were predicted using Barrnap v0.9 (https://github.com/tseemann/barrnap#barrnap) with Eukaryota database and categorized in 5S, 5.8S, 18S, and 28S rRNAs. INFERNAL v1.1.4^70^ was used to search for miRNAs and snoRNAs based on Rfam14.9 database^71^ **(Supplementary Table 12)**.

### Functional annotation and enrichments

Ortholog genes were identified by searching NCBI BLAST^72^ with default parameters from the NR database using default parameters. The top BLAST was reported for each gene. To annotate the transcripts for gene ontology (GO) terms and protein family (Pfam) terms, BLAST results were imported to OmicsBox software v2.2.4 (BioBam, Spain). The predicted protein sequences were searched against Pfam, ProDom, ProSiteProfiles by InterProScan in OmicsBox to retrieve conserved domains/motifs and corresponding GO terms using default parameters. Omicsbox (Blast2Go suite) functional annotation pipeline was followed, with mapped and annotated GO terms being selected for analysis^73^. The Kyoto Encyclopedia of Genes and Genome (KEGG) orthology (KO) and pathways were identified using the Automatic Annotation Server (KAAS; v2.1; http://www.genome.jp/kegg/kaas/) with bi-directional best hit (BBH) and BLAST as search program against closely related organisms, mallow, rose, mustard, and papaya. Carbohydrate-active enzymes were annotated from the proteome with the trinotate software (v3.2.2) using the hmmscan function against the Carbohydrate Active Enzymes (CAZy) database (http://www.cazy.org/) with default settings^74^. The plant Transcription factor & Protein Kinase Identifier and Classifier (iTAK) online tool v1.6 (http://itak.feilab.net/cgi-bin/itak/online_itak.cgi) was used to identify transcription factor families with default settings. Enrichment analysis for all functional annotations was performed using a Fisher’s test, and the resulting P-values were adjusted with the Benjamini and Hochberg method^75^. Enrichments with *p_adj_*<0.05 were considered significant.

### RNA extraction, cDNA library preparation, RNA sequencing, and sequencing data processing of nut tissues for spatiotemporal study

Fruit tissues from 15 of the 24 weeks assessed for physiological data were dissected into the hull, shell, and kernel and were each flash-frozen in liquid nitrogen on the day of sampling. For the kernel, the maternal seed coat was removed such that only the embryo was frozen. Four replicates composed of 12 fruits from three separate trees, from the same fruits used for physiological measurements, were frozen from the same fruits used for physiological measurements. Frozen tissues were ground into a fine powder with the Retsch Mixer Mill MM 400 (Verder Scientific, Netherlands). RNA extractions were performed on samplings from 865-2,564 GDD for hull tissues, 865-2139 GDD for shell tissues, and 1,106-2,564 GDD for kernel tissues. One gram of ground tissue was used for RNA extraction as described in Blanco-Ulate *et al*. 2013^76^. RNA concentrations were quantified with Nanodrop One Spectrophotometer (Thermo Scientific, United States) and Qubit 3 (Invitrogen, United States). RNA integrity was then assessed on an agarose gel.

RNA was extracted and sequenced from at least three biological replicates per time point and tissue. cDNA libraries were prepared with Illumina TruSeq RNA Sample Preparation Kit v.2 (Illumina, United States) from the extracted RNA. The quality of the barcoded cDNA libraries was assessed with the High Sensitivity DNA Analysis Kit in the Agilent 2100 Bioanalyzer (Agilent Technologies, United States) and then sequenced (150 bp paired-end reads) on the Illumina HiSeq X Ten platform by IDseq Inc. (Davis, CA, United States).

Raw reads were trimmed for quality and adapter sequences using Trimmomatic v0.39^57^ with the following parameters: maximum seed mismatches = 2, palindrome clip threshold = 30, simple clip threshold = 10, minimum leading quality = 3, minimum trailing quality = 3, window size = 4, required quality = 15, and minimum length = 36. Trimmed reads were then mapped using Bowtie2 v2.3.4.1^77^ to the predicted pistachio 37,955 protein sequences, producing an average of 76.41% mapping percentage across all samples (**Supplementary Table 14**). Count matrices were made from the Bowtie2 results using sam2counts.py v0.913 (https://github.com/vsbuffalo/sam2counts/blob/master/sam2counts.py). Read counts were normalized with the Bioconductor package DESeq2 in R^78^. Gene expression visualizations used the median of ratios normalization (DESeq2) for cross-sample comparison (**Supplementary Table 9**). rRNA genes identified (see *“Genome Annotation”* section above) were excluded from the count matrix for gene expression analysis.

### Gene co-expression analysis

Count matrices were normalized for gene co-expression analysis with the variance stabilizing transformation (vst) function from the DESeq2 package in R. The normalized reads were then used for weighted gene co-expression network analysis with the WGCNA package in R^79^. The WGCNA was conducted for each tissue separately. Transcripts that had a count of <10 reads in 90% of each tissue’s samples were removed and then checked by the goodSamplesGenes function. The dendrogram plots were made to identify and remove sample outliers. One kernel sample was removed with a height of more than 200 (h>200), and four hull samples were removed with a height of more than 120 (h>120). The network construction was done step by step. The soft thresholding power was selected to obtain the approximate scale-free topology, and adjacency was calculated with power equals selected soft power and type is signed type. TOM was calculated with signed type. The initial module eigengenes were detected by default hierarchical clustering function with default parameters other than the appropriate minModuleSize selected for each tissue, then final module eigengenes were obtained by merging close modules with cutHeight = 0.25. The module eigengenes values were then used to identify modules significantly associated with measured physiological traits by calculating correlations using Pearson’s product-moment correlation coefficient. The intramodular connectivity was assessed using the edge file output based on TOM.

### Macronutrient measurements

Non-structural carbohydrates (NSC) were analyzed in pistachio kernels. Frozen ground tissue was utilized as described above in “*Sample collection for physiological data, RNAseq experiments, and metabolic analysis.”* The concentration of NSC, including soluble sugars and starch, was analyzed as described in previous publications^80,81^, with some modifications. For soluble sugar extraction, 25 mg of fresh sample was mixed with 1 mL 0.2 M sodium acetate buffer, pH 5.5, and incubated at 70°C for 15 min. After centrifugation at 21,000 g for 10 min, the supernatant was diluted in bi-distilled water (1:10, v/v). Soluble sugars were then quantified by adding 0.1 % anthrone in 98 % sulfuric acid (m/v), incubating at 100 °C for 20 min, cooling at room temperature for 10 min, and reading absorbance at 620 nm in a photometer (Multiskan FC, Thermo Scientific, United States) and using a glucose standard curve to compare the colorimetric response of the samples. For starch extraction, the remaining buffer and pellet were incubated at 100 °C for 10 min and cooled for 20 min. Subsequently, 0.7 U of amylase and 7 U of amyloglucosidase (Sigma-Aldrich, United States) were added. The samples were stirred in a rotary incubator at 37°C for 4 h. Samples were then centrifuged at 21,000 g for 10 min, and the supernatant was diluted in bi-distilled water (1:10, v/v). Soluble sugars were quantified as described before. Starch concentration was determined by subtracting pre-starch digestion soluble sugar concentration from total (pre- and post-starch digestion) soluble sugar concentration. Three repetitions per sample were analyzed, and the results were averaged.

Protein content was measured as crude protein with AOAC Official Method 990.03 at the UC Davis Analytical Lab, University of California, Davis, United States. Total crude protein was calculated from the nitrogen content with the protein factor 6.25 applied to the nitrogen result.

### Volatile profiling

Volatiles were measured following the methodology described in Polari and colleagues^38^. Briefly, fresh pistachio kernels were ground to a paste in a mortar, and a sample (1.5 ± 0.1 g) was spiked with dodecane (2.5 mg/kg nutmeat). Deionized water (3 mL) was added, and the vial was sealed with a PTFE/silicon septum (Supelco, United States). After 15 min at 50℃, a solid-phase microextraction (SPME) fiber (DVB/CAR/PDMS, Sigma-Aldrich, United States) was exposed to the sample headspace for 45 min for volatile extraction. The volatile compounds analysis was performed with a Varian 450-GC equipped with a Varian 220-MS ion trap (Agilent Technologies, United States). A DB-5MS capillary column (30 m · 0.25 mm · 0.25 mm, Agilent Technologies, United States) was used for compound separations. After sampling, the fiber was thermally desorbed in the GC injector for 5 min at 260℃. Helium was the carrier gas at a flow rate of 1 mL/min. The oven temperature started at 40℃ during the 5-min desorption process and was ramped at 3.5℃/min to 120℃, then ramped at 10℃/min to 220℃ and held for 10 min. An ionization energy of 70 eV was adopted, and the ions were analyzed in the m/z range from 40 to 400. The data were recorded and analyzed using MS Workstation v. 6.9.3 (Agilent Technologies, United States) software. Volatile compounds were identified by their mass spectra and Kovats retention index (KI). Results were expressed as nanograms of dodecane per kilogram of sample.

### Phenolic measurements

Three biological replicates were analyzed for three sampling dates: August 25 (2,162 GDD), August 31 (2,279 GDD), and September 7 (2,402 GDD). Total phenolics were extracted from the kernels following the protocol detailed in Grilo and colleagues^79^. Phenolic extracts were membrane-filtered with 0.45 µm cellulose and subjected to HPLC-DAD analysis as described by Erşan and colleagues^80^. HPLC-DAD model (Agilent G4212-60008, serial number-DEBAF01604, Agilent Technologies, United States) was used with the following solvents: HPLC grade water (99%, Solvent A) and HPLC grade methanol (99%, Solvent B), with each solvent containing 1% (v/v) formic acid. The column was an Eclipse Plus C18 column (250 mm x 4.6 mm, 5 µm; Agilent Technologies, USA) with a security guard ultra C18 guard column (4.6 mm x 2 mm) of the same material. The gradient included: isocratic at 2% B for 10 min, then 2 to 37% B in 27 min, isocratic at 37% B for 5 min, then from 37 to 40 % B in 18 min, from 40 to 60% B in 10 min, from 60 to 100% B in 20 min, isocratic at 100 % B for 14 min, then from 100 to 2% B in 1 min, and isocratic at 2% B for 7 min. The column temperature was 35°C, and the total run time was 112 min at a flow rate of 1 mL/min and injection volume of 3 µL. The calibration curves with the external standards were obtained using concentration (mg L^−1^) with respect to the area obtained from the integration performed at 280 nm (gallotannins), 310 nm (anacardic acids), and 350 nm (flavonols). Results were expressed as mg/g of kernel dry weight.

### Fatty acid profiling

Fatty acid profiles were determined according to Polari and colleagues^38^, as described. Briefly, the sample (0.010 ± 0.001 g) was weighed and dissolved in toluene (0.4 mL). Methanol (3 mL) and methanol/HCl (0.6 mL, 80:20, v/v) were added. The vial was heated at 80℃ for 1 h. Hexane (1.5 mL) and deionized water (1 mL) were added. Anhydrous sodium sulfate (0.5 g) was added to the decanted upper hexane layer containing the methyl esters. GC analysis was conducted on a Varian 450-GC (Agilent Technologies, United States) with a flame ionization detector. Helium was the carrier gas at a flow rate of 1.5 mL/min. Fatty acid methyl esters (1 ml) were injected onto a DB-23 capillary column (60 m · 0.25 mm · 0.25 mm (Agilent Technologies, United States). A mix of FAME standards was used as references for peak identification by retention times; relative fatty acid proportions were determined by peak area normalization. Individual fatty acids were expressed as the percentage of total fatty acids.

### Hormone measurements

Hull, shell, and kernel tissues were taken from the 2019 samples used for physiological and transcription analysis (see *Sample collection for physiological data, RNA-seq experiments, and metabolic analysis*). Sample analysis was provided by Metware Biotechnology Inc (Boston, MA, USA). Briefly, 15 mg of frozen plant sample was dissolved in 1 mL methanol/water/formic acid (15:4:1, v/v/v). 10 μL internal standard mixed solution (100 ng/mL) was added into the extract as internal standards (IS) for the quantification. The mixture was vortexed for 10 minutes, followed by centrifugation for 5 min (12000 r/min, and 4°C). The supernatant was transferred to clean plastic microtubes and dried by evaporation. The samples were reconstituted in 100 μL 80% methanol (v/v), and filtered through a membrane filter (0.22 μm, Anpel) before LC-MS/MS analysis^84–86^. The sample extracts were analyzed using a UPLC-ESI-MS/MS system (UPLC, ExionLC™ AD, https://sciex.com/; MS, Applied Biosystems 6500 Triple Quadrupole, https://sciex.com/). UPLC Conditions were as follows: Column: Waters ACQUITY UPLC HSS T3 C18 (100 mm×2.1 mm i.d.,1.8 µm); solvent system: water (0.04% acetic acid):acetonitrile (0.04% acetic acid); gradient program: 95:5 v/v at 1 min, 5:95 v/v at 8 min, hold for 1 min, 95:5 v/v at 9 min; hold for 3min; flow rate: 0.35 mL/min; temperature: 40°C; injection volume: 2 μL.

The mass spectrum conditions were as follows: Triple quadrupole (QQQ) scans were acquired on a triple quadrupole mass spectrometer, 6500+ LC-MS/MS System, equipped with an ESI Turbo Ion-Spray interface, operating in both positive and negative ion mode and controlled by Analyst 1.6.3 software (Sciex, United States). The ESI source operation parameters were as follows: ion source, ESI+/-; source temperature 550℃; ion spray voltage (IS) 5500 V (Positive) - 4500 V (Negative); curtain gas (CUR) was set at 35 psi, respectively. Phytohormones were analyzed using scheduled multiple reaction monitoring (MRM). Data acquisitions were performed using Analyst 1.6.3 software (Sciex, United States). Multiquant 3.0.3 software (Sciex, United States) was used to quantify all metabolites. Mass spectrometer parameters including the declustering potentials (DP) and collision energies (CE) for individual MRM transitions were done with further DP and CE optimization. A specific set of MRM transitions was monitored for each period according to the metabolites eluted within this period^84,87,88^.

### PvNFYB-LEC1 and PvAP2-WRI1 genes and predicted binding motif analysis

The WRINKLED1 (WRI1) (accession: Q6X5Y6) and LEAFY COTYLEDON1 (LEC1) gene (accession: Q9SFD8) of *Arabidopsis thaliana* from the UniProt (https://www.uniprot.org/) were used to identify homologs of AtWRI1 and AtLEC1 in ‘Kerman’ and 14 other taxa **(Supplementary Table 15)**. *A*. *thaliana* WRI1 was used as a query to BLAST against filtered proteomes of 15 taxa and BLAST hits with similar size to *A*. *thaliana* WRI1 genes were analyzed for the conserveness of two AP2/EREBP DNA binding domains. For LEC1, because the sequences outside the central conserved region showed high divergence between AtLEC1, BnLEC1(*Brassica napus*, EU371726), and GmLEC1 (*Glycine max*, Glyma.07G268100), 103 aa sequences in the highly conserved region in AtLEC1 were used for the BLAST search with e-value 1e-10. BLAST hits were selected with the highest similarity, and genes were extracted from the proteomes as possible LEC1 genes. Amino acid sequence alignments of all 16 WRI1 and LEC1 genes were performed using MAFFT v7.505^89^. The 3D structure of all WRI1 and LEC1 proteins was predicted using ColabFold ^90^, which utilizes AlphaFold2 ^91^ and visualized in USCF ChimeraX^68^ with coloring based on the predicted local distance difference test (pLDDT) scores. To search for binding sites of WRI1 and LEC1 genes in the ‘Kerman’ genome, the consensus sequence (CNTNG(N_7_)CG) of AW-box binding site for WRI ^27,28,92^ and ‘CCAAT’ for LEC1 were examined in nucleotide sequences 1,500 bp upstream of the translational initiation site (TIS) of gene models in ‘Kerman’ genome with a focus on genes associated with fatty acid biosynthesis and the K-III-1 gene module. The consensus sequences of AW-box binding sites were visualized using WebLogo^93^. The distribution of binding sites of WRI1 was calculated to examine the relative distance from the TISs in genes of fatty acid biosynthesis and the K-III-1 gene module compared to other genes with the AW-box binding motif.

## Supplementary Material

**Supplementary Figure 1.** Nut physiological patterns are consistent across years and locations. Stages were confirmed by assessing, a, whole nut and kernel area growth, b, dry weight (g) of whole nut and kernel, color changes in the kernel, c, and hull, d, measured in the L*a*b* color space, (L*, or lightness, a* or redness, b* or yellowness), e, texture changes in the hull, shell, and kernel (kg of Force) in three years and orchard locations. For trait specific dynamics across the season and environmental conditions, we fitted linear and linear-mixed polynomial models for each physiological trait as a function of accumulated heat (GDD). To test if the behavior of these traits was consistent across different environmental conditions we fitted models for multi-years data and found that nut size and weight could be accurately modeled, while color and texture changes were more subject to environmental variation.

**Supplementary Figure 2.** Estimation of genome size, heterozygosity, and repetitiveness with *k*-mer frequency using jellyfish and GenomeScope. K-mer size was set to 21 with coverage cutoff at 1e6 to contain repetitive regions in the estimation. len, haploid length; uniq, percentage of unique (non-repetitive) sequence; het, percentage of heterozygosity; err, percentage of sequencing error rate; dup, rate of mean read duplication.

**Supplementary Figure 3.** Repeat analysis in P. vera ‘Kerman’ genome assembly. A) The amount of total repetitive sequences in genome assemblies of P. vera ‘Kerman’, P. vera ‘Batoury’, Mango, Sweet orange, and Trifoliate orange. B) Proportion of different transposable element (TE) types in ‘Kerman’ genome assembly.

**Supplementary Figure 4.** Macro-synteny comparison between genome assemblies of P. vera ‘Kerman’ and three other cultivars (‘Batoury’, ‘Siirt’, and ‘Bagyolu’). Lower percent identity regions were filtered out in dot-plots on the right side compared to the default settings on the left side.

**Supplementary Figure 5.** The number of unique and overlapping transcripts expressed in five different tissue types. All five tissue types share 9753 transcripts and 450, 398, 376, 326, and 265 transcripts were unique in developing buds, leaves, dormant buds, fruits, and flowers, respectively.

**Supplementary Figure 6.** Weighted gene co-expression network analysis (WGCNA) was performed for genes expression across all time points for each tissue type. The resulting groups of co-expressed genes (modules) were categorized by their eigengene value patterns (a) to the stages in development that changes were occurring (labeled by color). These patterns were correlated to the physiological patterns observed in sample (Figure 1, Supplementary Table 1) to produce trait-module relationships for each tissue type (b). The heatmap color indicates the type and strength of the correlation (R^2^), either negative or positive. Values in the squares indicate the R^2^ value (top) and the significance of that correlation (p-value, bottom).

**Supplementary Figure 7.** Alignment of the protein sequences of WRINKLED1 (WRI1) genes of 16 representative angiosperm species ordered based on phylogenetic relationships with A. trichopoda as an outgroup. AP2/EREBP DNA binding domains in ‘Kerman’ WRI1 are indicated with red annotations. The blue and purple annotations represent the VYL domain encoded by 9 bp exon 3 and the sites phosphorylated by KIN10 (T70 and S166), respectively. Folding confidence level are indicated in different colors.

**Supplementary Figure 8.** Alignment of the protein sequences of LEAFY COTYLEDON1 (LEC1) genes of 16 representative angiosperm species ordered based on phylogenetic relationships with A. trichopoda as an outgroup. DNA binding domain in ‘Kerman’ LEC1 is indicated with red annotation. Folding confidence level are indicated in different colors.

**Supplementary Figure 9.** Workflow for PacBio HiFi sequencing, transcript discovery, and *de novo* genome assembly and scaffolding from sample collections in the field to chromosome-scale genome.

**Supplementary Figure 10.** Genome annotation pipeline using both RNA- and Iso-seq data as extrinsic hints. Repetitive regions were masked using EDTA and ProtExcluder. After automated training with Braker2, ab initio gene prediction was performed using Augustus and EvidenceModeler followed by UTR and isoform variants updates with PASApipeline.

**Supplementary Table 1.** Summary of physiological data collected across different locations and harvest years (2019, 2020, 2021). Average, standard deviation (St.Dev.) and number of nuts (n) used for analysis are shown for each harvest date and cooresponding growing degree days (GDD) expressed in °C. Cells with a dash indicate no data collection at that time point.

**Supplementary Table 2.** Summary of physiological growth models. For trait specific dynamics across the season and environmental conditions, we fitted linear and linear-mixed polynomial models for each physiological trait as a function of accumulated heat (GDD). To test if the behavior of these traits was consistent across different environmental conditions we fitted models for multi-years data and found that nut size and weight could be accurately modeled, while color and texture changes were more subject to environmental variation. Data summarized in **Supplementary Table 1** was used to develop the models. The models for 2019 are represented in **Fig. 1**, while the models combining all years of data are shown in **Supplementary** Figure 1. Data was transformed where indicated.

**Supplementary Table 3.** Statistics of PacBio HiFi sequencing and Dovetail genomics Omni-C data

**Supplementary Table 4.** Statistics of genome assembly and annotation of the *P*. *vera* ‘Kerman’.

**Supplementary Table 5.** The summary of BUSCO (Benchmarking Universal Single-Copy Orthologs) assessment of ‘Kerman’ primary contig assembly and 15 chromosomes.

**Supplementary Table 6.** Summary statistics of repeat content in ‘Kerman’ primary contig assembly.

**Supplementary Table 7.** Statistics of the number of transcripts in different tissue types.

**Supplementary Table 8.** The summary of BUSCO (Benchmarking Universal Single-Copy Orthologs) assessment of gene annotation of ‘Kerman’ primary contig assembly and 15 chromosomes.

**Supplementary Table 9.** *P. vera* ‘Kerman’ transcriptome functional annotations and gene expression data. The average median of ratios normalized reads were calculated with the DESeq2 package (Love et. al 2014) with functional annotations and weighted gene coexpression network analysis (WGCNA) module assignment (Langfelder and Horvath 2008) for the three tissues (hull, shell, kernel) across three stages (Stage II, Stage III, Stage IV). The time points (expressed in growing degree days (GDD, °C) WGCNA was performed on expression data from each tissue across all time points. Genes with a minimum of 10 reads expression across the all timepoints were assigned a module. KEGG = Kyoto Encyclopedia of Genes and Genomes. iTAK = Transcription factor & Protein Kinase Identifier and Classifier. CAZy= Carbohydrate Active Enzymes Database. GO= Gene Ontology.

**Supplementary Table 10. Enrichments of functional terms.** Functional enrichments in Kyoto Encyclopedia of Genes and Genomes (KEGG) and iTAK Plant Transcription Factors and Protein Kinases among gene co-expression modules for each tissue (kernel, hull, shell). Adjusted p-values are shown for each module. Red highlighted p-values are significant (Padj<0.05) as determined by Fisher’s test adjusted bwith the Benjamini and Hochberg method. Dashes indicate no significance or no data. Only functional annotations with one or more significant modules are shown. Enrichment were run for the terms in the column ‘Pathway/Gene Family’. The column ‘Heirarchical Term’ corresponds to the category of the pathway/gene family in the respective functional ontology database. Hormone terms were curated from KEGG.

**Supplementary Table 11.** Summary of Metabolite Data. Metabolite data is summarized for volatiles, fatty acids, and phenolic compounds acorss sampling dates for 2019 and 2020 seasons. The corresponding growing degree days (GDD, degrees C) for each sampling date are indicated. Average, standard deviation (St.Dev.) and number of samples (n) analyzed are summarized.

**Supplemental Table 12.** Hormone measurements were taken at one timepoint at the end of Stage III on August 13, 2019 corresponding to 1881 growing degree days (GDD, degrees C). Abscisic acid (ABA), jasomonic acid (JA) and the bioactive form JA-ILE were measured. Average, standard deviation (St.Dev.) and number of samples (n) are summarized.

**Supplementary Table 13.** The statistics of non-coding RNA in *Pistacia vera* ‘Kerman’ genome

**Supplementary Table 14.** RNA-sequencing read mapping summary for pistachio hull, shell, and kernel tissue collected at each time point (growing degree days, GDD) in each stage. Parsed reads are those that passed through quality and adapter trimming.

**Supplementary Table 15.** List of angiosperm taxa used in comparative genomic analysis

## Data availability

All data included in this study can be found under the BioProject accession number PRJNA1114109. Under this, all the raw genome sequencing data have been deposited at NCBI (https://www.ncbi.nlm.nih.gov/), with the BioProject accession number PRJNA1049825 (PacBio HiFi, Omni-C, and Iso-Seq reads). The genome assembly, annotation, and 3D structure of gene models are available at the Pistachiomics database hosted by UC Davis (https://pistachiomics.sf.ucdavis.edu/). The nut transcriptomic data discussed in this publication have been deposited in NCBI’s Gene Expression Omnibus ^94^ with the BioProject accession number PRJNA1110275 and are accessible through GEO Series accession number GSE267225 (https://www.ncbi.nlm.nih.gov/geo/query/acc.cgi?acc=GSE267225).

## Acknowledgments

The authors would like to acknowledge and thank Caio Cattai de Andrade for sampling pistachios from the 2019 growing season; Adrian O. Sbodio for sampling nuts from the 2020 season and processing the nut samples in 2020-2021; Jose G. Barquero-Jackson for sampling nuts from the 2021 growing season; Paula Guzman Delgado for measuring starch and carbohydrates in pistachio kernels, Pedro Bello for technical support in capturing images from Figure 1d.; and collaborators Joseph Coelho and Ian Humrick (Maricopa Orchards), Joey Thomas and John Thomas (Dewey Farms) for providing access to their pistachio orchards for sampling.

## Author Contributions

**Author list:** Jaclyn A. Adaskaveg^1+^(**JAA**), Chaehee Lee^1+^(**CL**), Yiduo Wei^1^ (**YW**), Fangyi Wang^1^(**FW**), Filipa S. Grilo^2,3^(**FSG**), Saskia D. Mesquida-Pesci^1^(**SDM-P**), Matthew Davis^1^(**MD**), Selina C. Wang^3^(**SCW**), Giulia Marino^1^(**GMa**), Louise Ferguson^1^(**LF**), Patrick J Brown^1^(**PJB**), Georgia Drakakaki^1^(**GD**), Adela Mena-Morales^4^(**AM-M**), Annalisa Marchese^5^(**AM**), Antonio Giovino^6^(**AG**), Esaú Martínez^4^(**EM**), Francesco Paolo Marra^5^(**FPM**), Lourdes Marchante Cuevas^3^(**LMC**), Luigi Cattivelli^7^(**LC**), Paolo Bagnaresi^7^(**PB**), Pablo Carbonell-Bejerano^8^(**PC-B**), Grey Monroe^1^*(**GMo**), Barbara Blanco-Ulate^1^*(**BB-U**)

**Conceptualization:** JAA, CL, GD, PJB, AM-M, AM, AG, EM, FPM, LMC, LC, PB, PC-B, GMo, BB-U

**Methodology:** JAA, CL, FSG, SDM-P, MD, SCW, GMa, LF, GMo, BB-U

**Formal Analysis:** JAA, CL, YW, FW, FSG, MD, GMo, BB-U

**Resources:** AM-M, AM, AG, EM, FPM, LMC, LC, PB, PC-B, GMo, BB-U, SW, GMa,

**Investigation:** JAA, YW, SDM-P, FSG,

**Data Curation:** JAA, CL, YW, MD, GMo

**Writing - Original Draft:** JAA, CL, GMo, BB-U

**Writing - Review & Editing:** All

**Visualization:** JAA, CL, YW, MD, GMo, BB-U

**Supervision:** JAA, CL, PCB, PJB, GMo, BB-U

**Project Administration:** PC-B, GMo, BB-U

**Funding Acquisition:** GMa, SCW, AM-M, AM, AG, EM, FPM, LMC, LC, PB, PC-B, GMo, PJB, BB-U

