## Supplementary Figures for "In a nutshell: pistachio genome and kernel development"

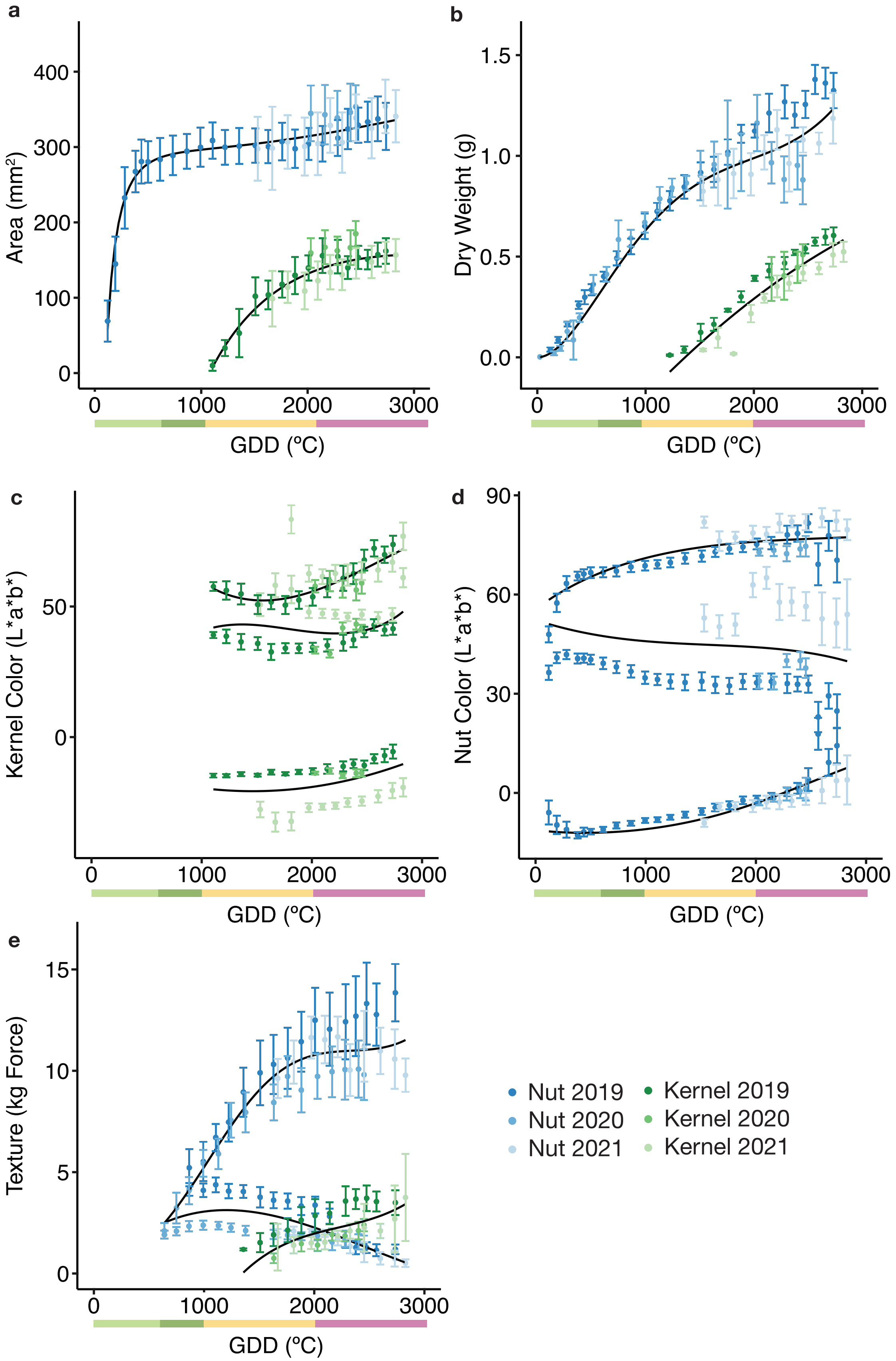


**Supplemental Figure 1. Nut physiological patterns are consistent across years and locations.** Stages were confirmed by assessing, a, whole nut and kernel area growth, b, dry weight (g) of whole nut and kernel, color changes in the kernel, c, and hull, d, measured in the L*a*b* color space, (L*, or lightness, a* or redness, b* or yellowness), e, texture changes in the hull, shell, and kernel (kg of Force) in three years and orchard locations. For trait specific dynamics across the season and environmental conditions, we fitted linear and linear-mixed polynomial models for each physiological trait as a function of accumulated heat (GDD). To test if the behavior of these traits was consistent across different environmental conditions we fitted models for multi-years data and found that nut size and weight could be accurately modeled, while color and texture changes were more subject to environmental variation.

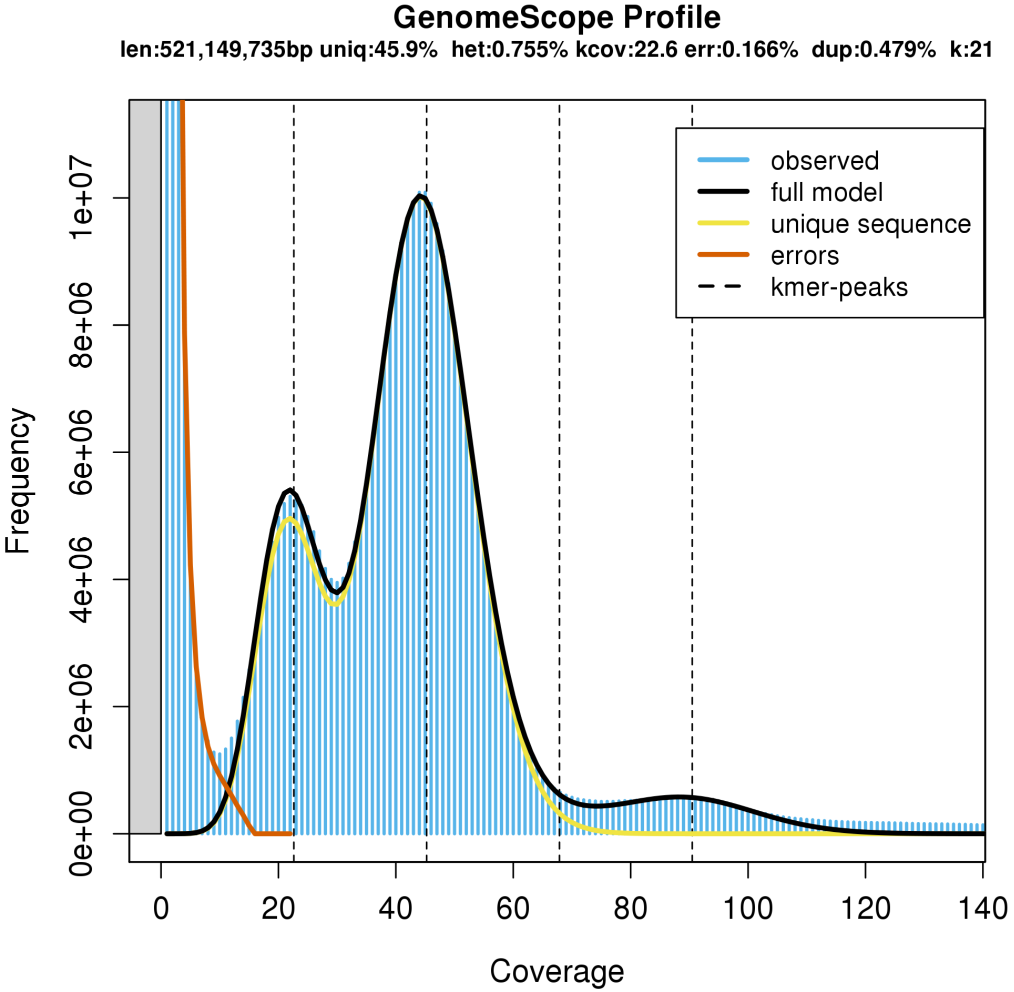


**Supplementary Figure 2. Estimation of genome size, heterozygosity, and repetitiveness with k-mer frequency using jellyfish and GenomeScope.** K-mer size was set to 21 with coverage cutoff at 1e6 to contain repetitive regions in the estimation. len, haploid length; uniq, percentage of unique (non-repetitive) sequence; het, percentage of heterozygosity; err, percentage of sequencing error rate; dup, rate of mean read duplication.

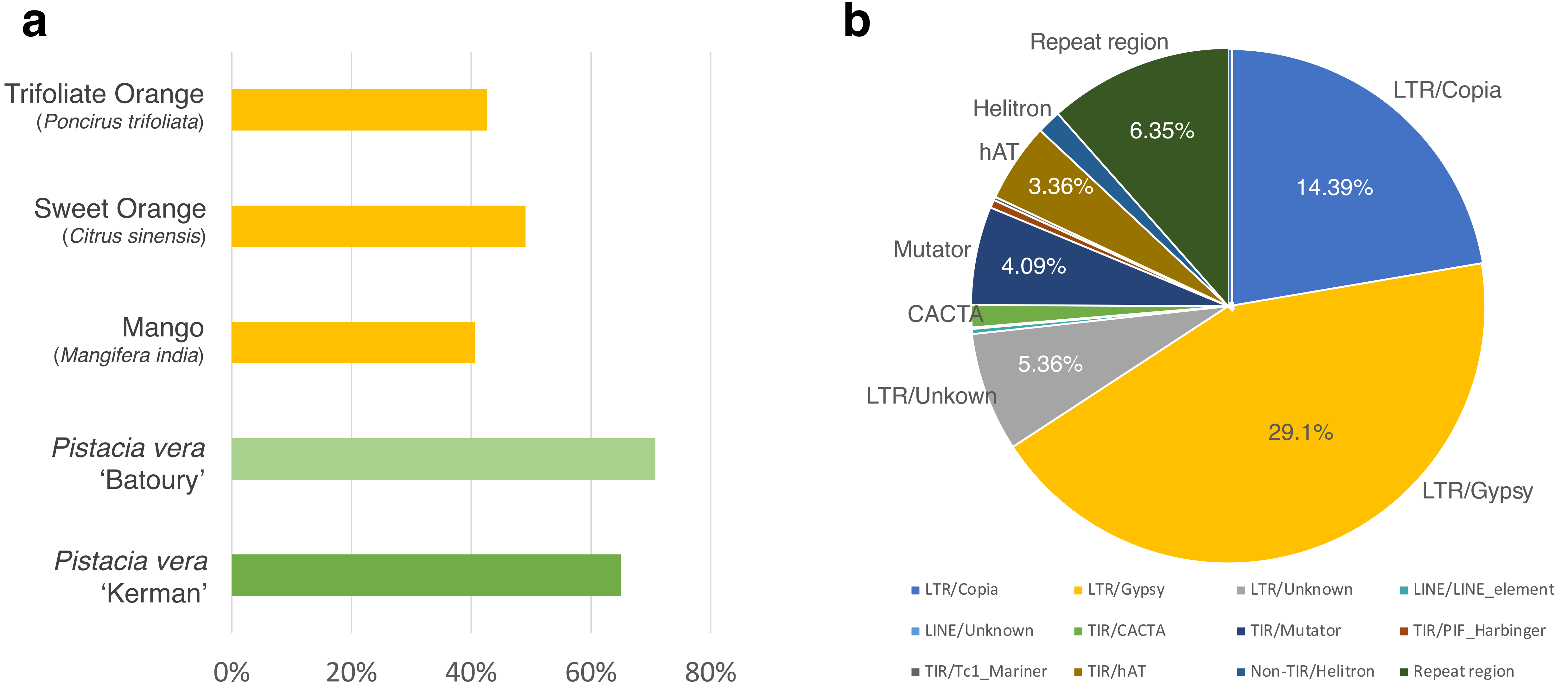


**Supplementary Figure 3. Repeat analysis in *P. vera* 'Kerman' genome assembly.** A) The amount of total repetitive sequences in genome assemblies of *P. vera* 'Kerman', *P. vera* 'Batoury', Mango, Sweet orange, and Trifoliate orange. B) Proportion of different transposable element (TE) types in 'Kerman' genome assembly.

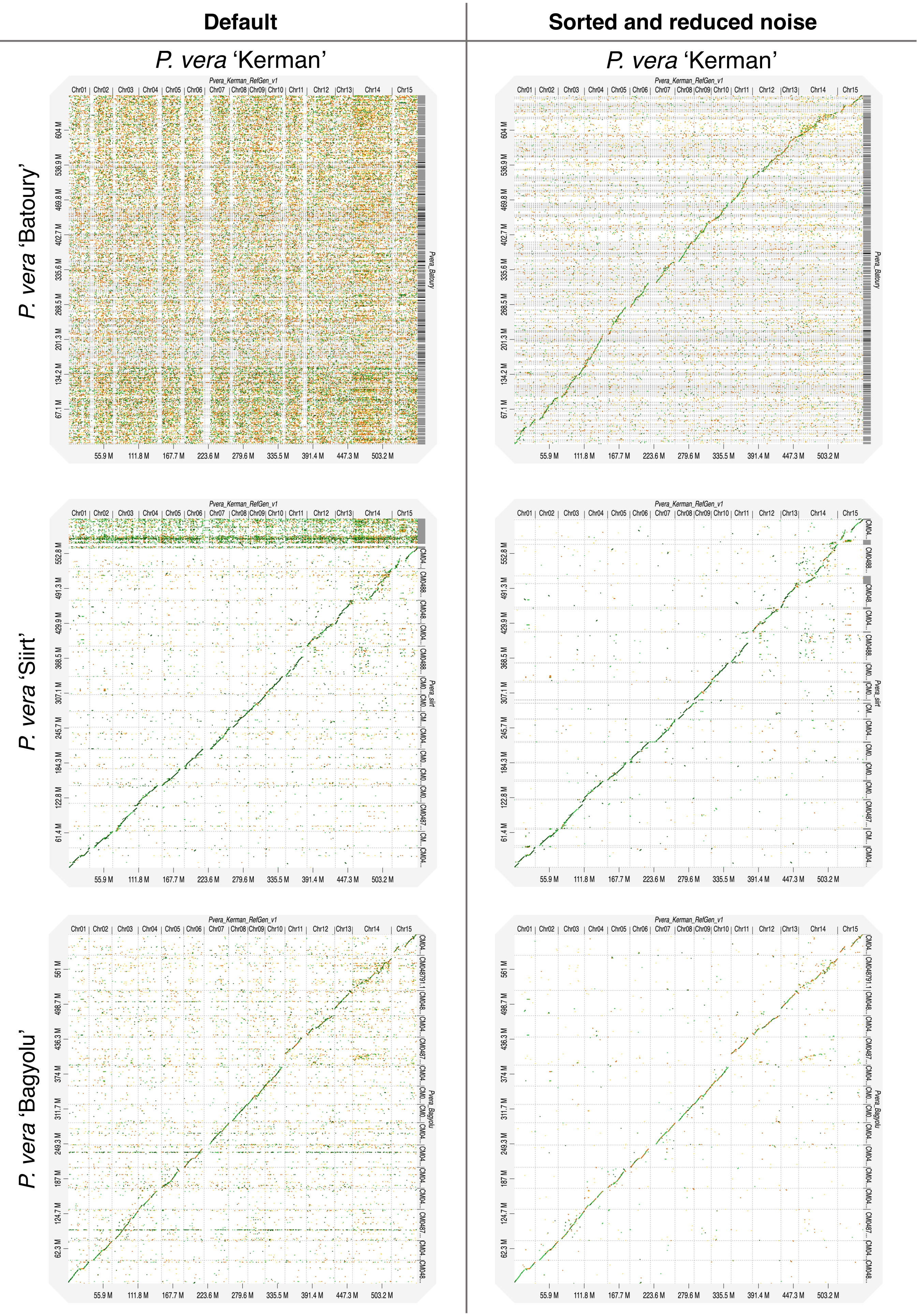


**Supplementary Figure 4. Macro-synteny comparison between genome assemblies of *P. vera* 'Kerman' and three other cultivars ('Batoury', 'Siirt', and 'Bagyolu').** Lower percent identity regions were filtered out in dot-plots on the right side compared to the default settings on the left side.

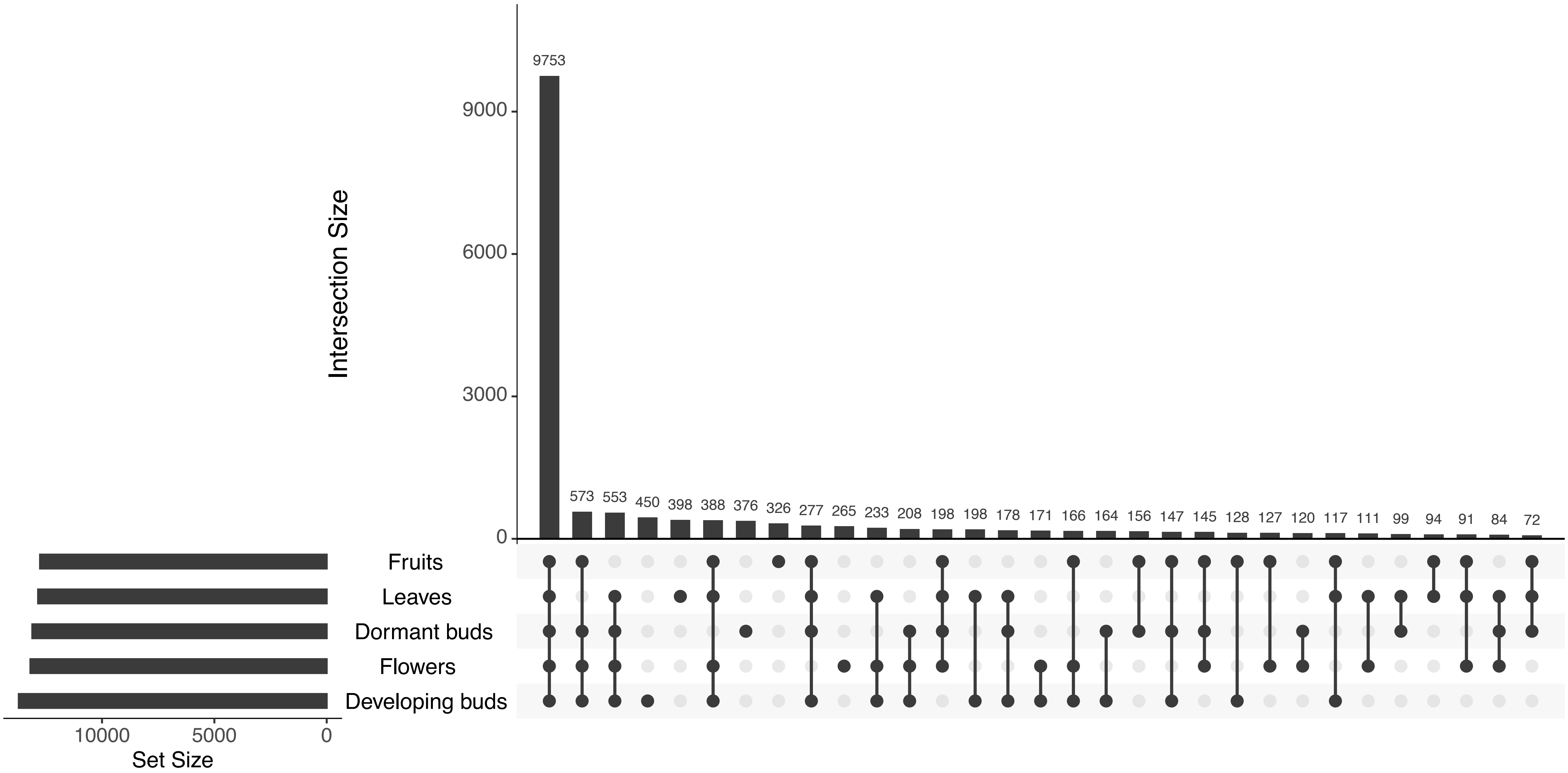


**Supplementary Figure 5. The number of unique and overlapping transcripts expressed in five different tissue types.** All five tissue types share 9753 transcripts and 450, 398, 376, 326, and 265 transcripts were unique in developing buds, leaves, dormant buds, fruits, and flowers, respectively.

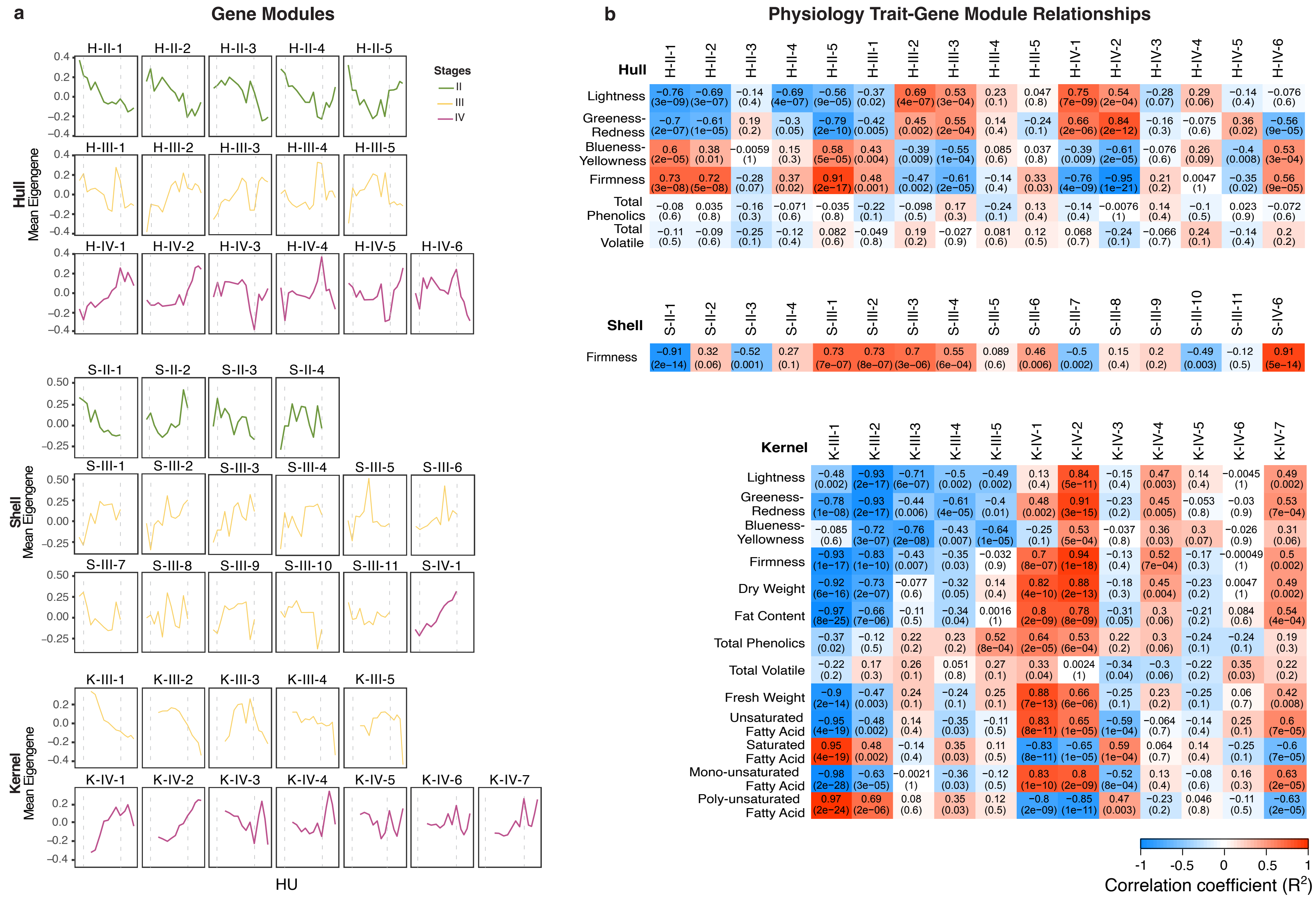


**Supplementary Figure 6. Weighted gene co-expression network analysis (WGCNA) was performed for genes expression across all time points for each tissue type.** The resulting groups of co-expressed genes (modules) were categorized by their eigengene value patterns (a) to the stages in development that changes were occurring (labeled by color). These patterns were correlated to the physiological patterns observed in sample (Figure 1, Supplementary Table 1) to produce trait-module relationships for each tissue type (b). The heatmap color indicates the type and strength of the correlation ($R^2$), either negative or positive. Values in the squares indicate the $R^2$ value (top) and the significance of that correlation (p-value, bottom).

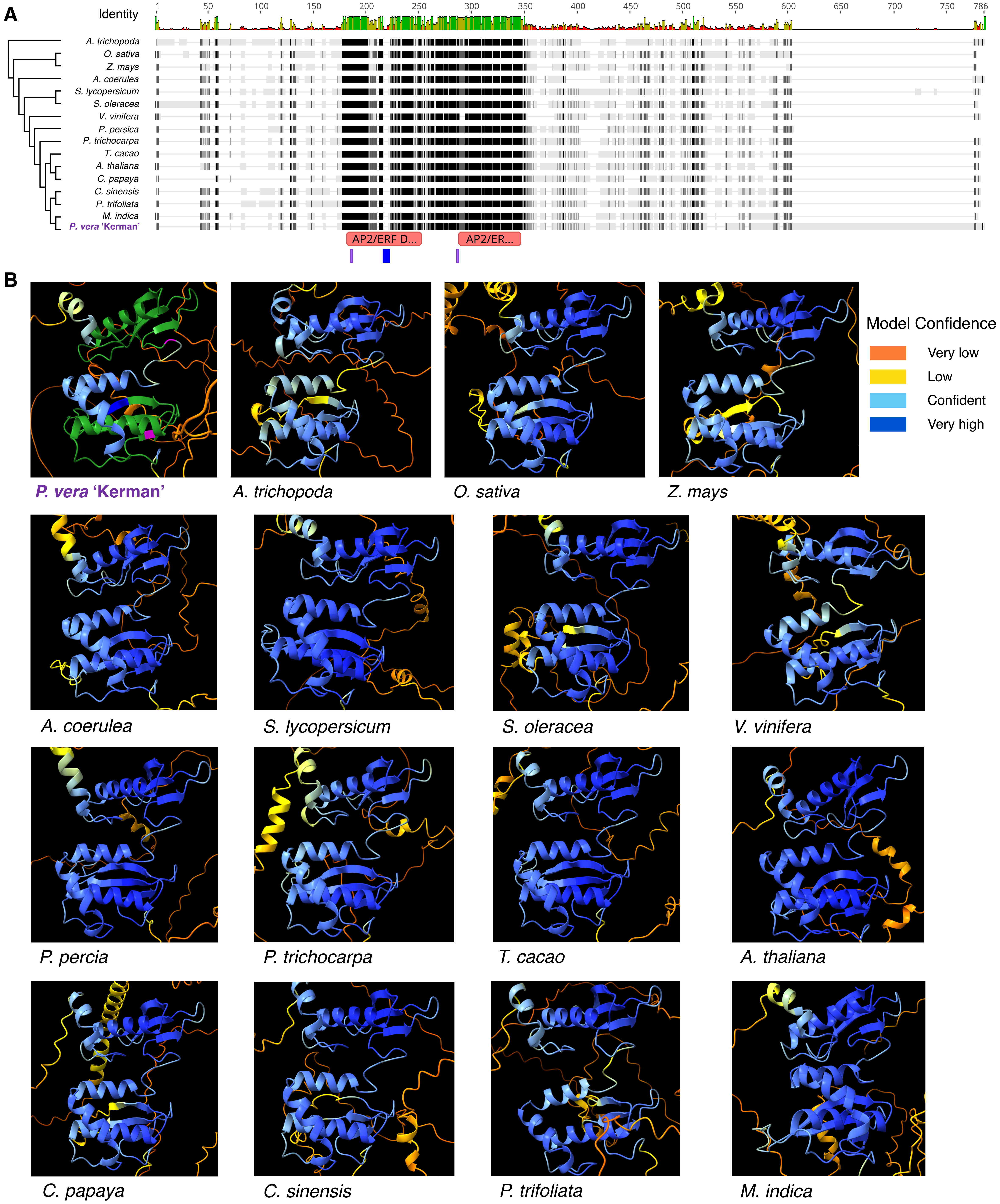


**Supplementary Figure 7. Alignment of the protein sequences of WRINKLED1 (WRI1) genes of 16 representative angiosperm species ordered based on phylogenetic relationships with *A. trichopoda* as an outgroup.** AP2/EREBP DNA binding domains in 'Kerman' WRI1 are indicated with red annotations. The blue and purple annotations represent the VYL domain encoded by 9 bp exon 3 and the sites phosphorylated by KIN10 (T70 and S166), respectively. Folding confidence level are indicated in different colors.

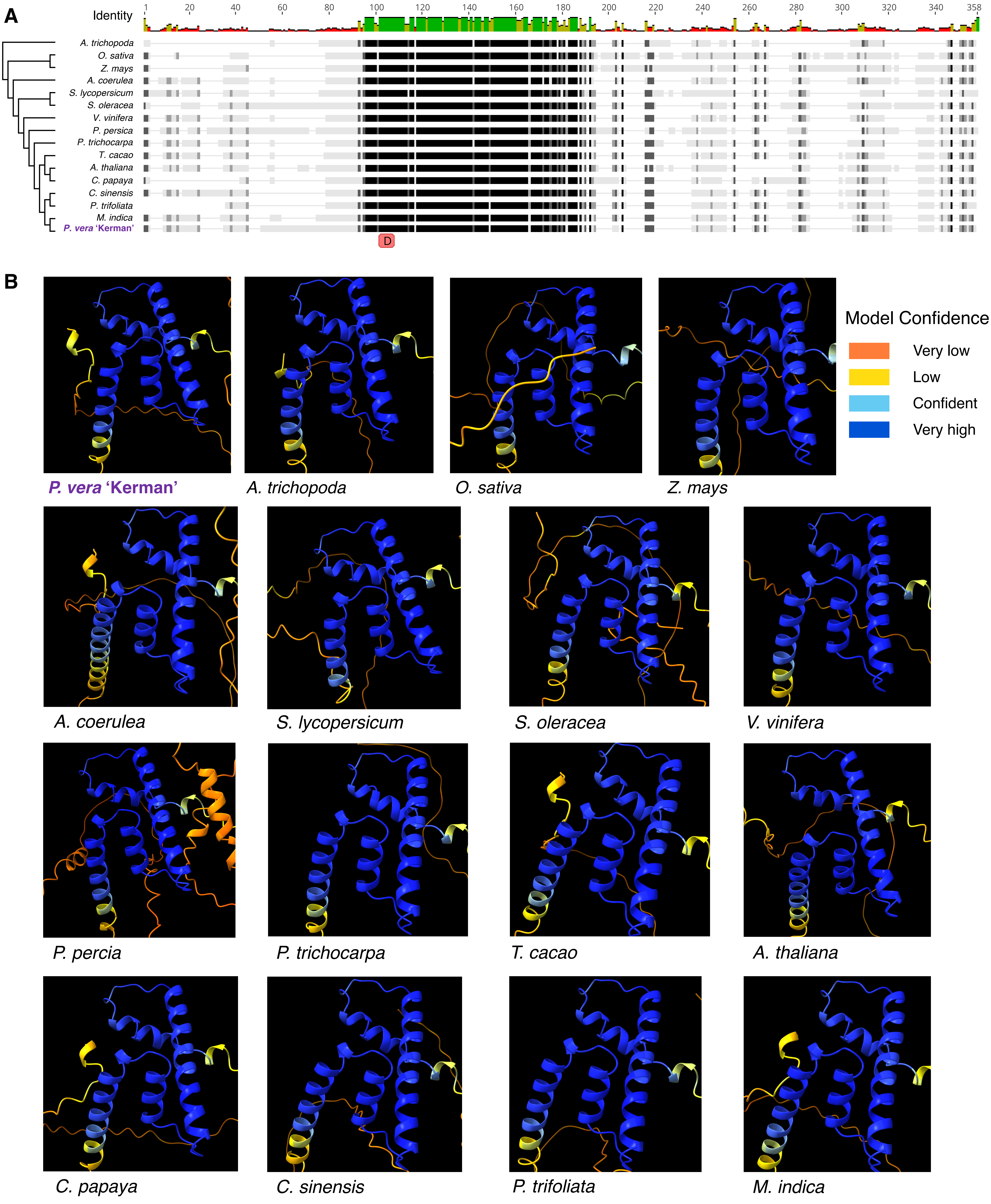


**Supplementary Figure 8. Alignment of the protein sequences of LEAFY COTYLEDON1 (LEC1) genes of 16 representative angiosperm species ordered based on phylogenetic relationships with *A. trichopoda* as an outgroup.** DNA binding domain in 'Kerman' LEC1 is indicated with red annotation. Folding confidence level are indicated in different colors.

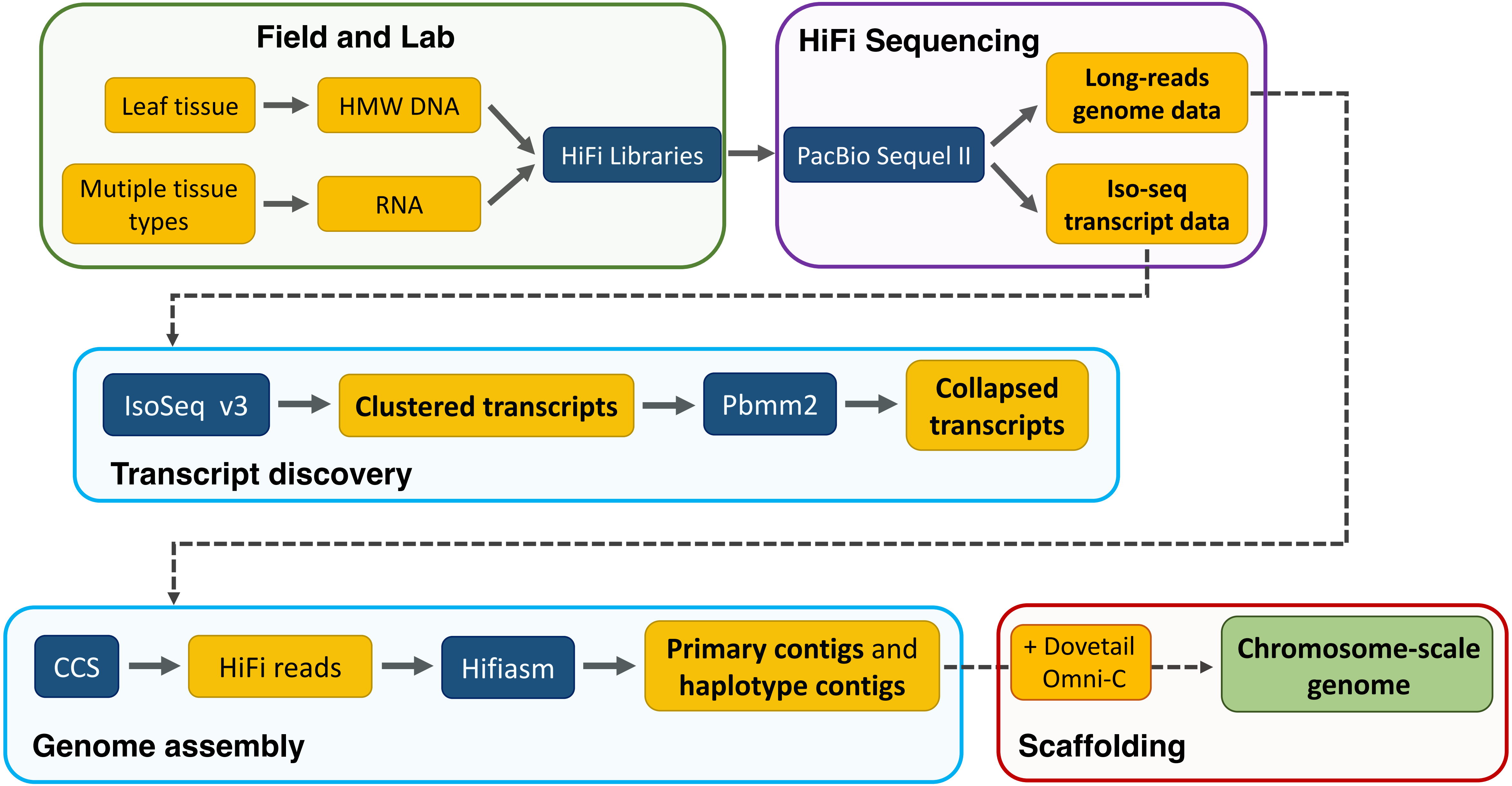


**Supplementary Figure 9. Workflow for PacBio HiFi sequencing, transcript discovery, and *de novo* genome assembly and scaffolding from sample collections in the field to chromosome-scale genome.**

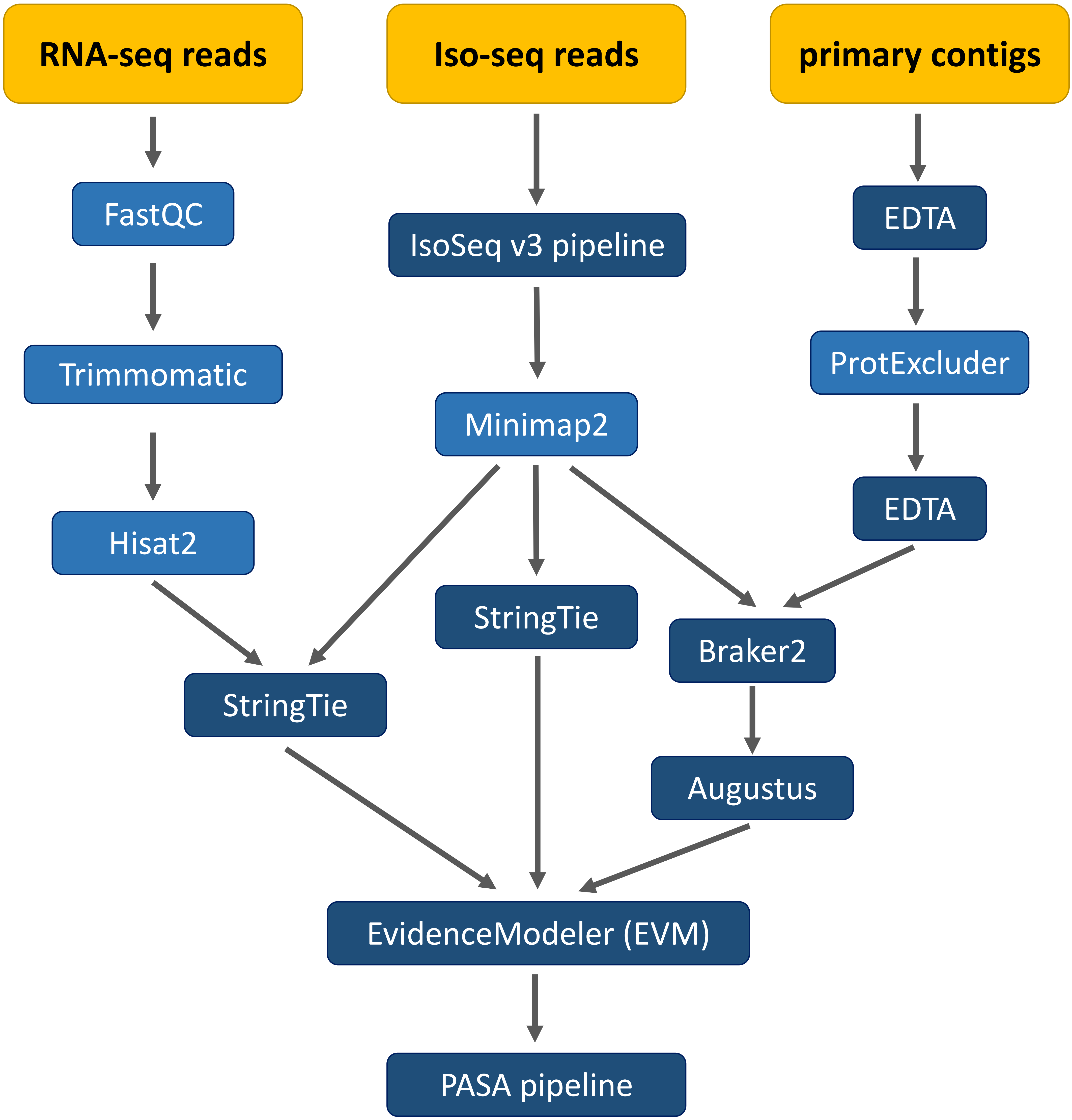


**Supplementary Figure 10. Genome annotation pipeline using both RNA- and Iso-seq data as extrinsic hints.** Repetitive regions were masked using EDTA and ProtExcluder. After automated training with Braker2, *ab initio* gene prediction was performed using Augustus and EvidenceModeler followed by UTR and isoform variants updates with PASApipeline.
